# Identification of Cellular Interactions in the Tumor Immune Microenvironment Underlying CD8 T Cell Exhaustion

**DOI:** 10.1101/2023.11.09.566384

**Authors:** Christopher Klocke, Amy Moran, Andrew Adey, Shannon McWeeney, Guanming Wu

**Affiliations:** Division of Bioinformatics and Computational Biomedicine, Department of Medical Informatics and Clinical Epidemiology, Oregon Health & Science University, Portland, OR, USA; Department of Cell, Developmental and Cancer Biology, Oregon Health & Science University, Portland, OR, USA; Knight Cancer Institute, Oregon Health and Science University, Portland, OR, USA; Department of Molecular and Medical Genetics, Oregon Health & Science University, Portland, OR, USA; Oregon Clinical and Translational Research Institute, Oregon Health & Science University, Portland, OR, USA

## Abstract

While immune checkpoint inhibitors show success in treating a subset of patients with certain late-stage cancers, these treatments fail in many other patients as a result of mechanisms that have yet to be fully characterized. The process of CD8 T cell exhaustion, by which T cells become dysfunctional in response to prolonged antigen exposure, has been implicated in immunotherapy resistance. Single-cell RNA sequencing (scRNA-seq) produces an abundance of data to analyze this process; however, due to the complexity of the process, contributions of other cell types to a process within a single cell type cannot be simply inferred. We constructed an analysis framework to first rank human skin tumor samples by degree of exhaustion in tumor-infiltrating CD8 T cells and then identify immune cell type-specific gene-regulatory network patterns significantly associated with T cell exhaustion. Using this framework, we further analyzed scRNA-seq data from human tumor and chronic viral infection samples to compare the T cell exhaustion process between these two contexts. In doing so, we identified transcription factor activity in the macrophages of both tissue types associated with this process. Our framework can be applied beyond the tumor immune microenvironment to any system involving cell-cell communication, facilitating insights into key biological processes that underpin the effective treatment of cancer and other complicated diseases.

## 1 Introduction

Cancer immunotherapies show great potential for treating patients with otherwise incurable disease. Over the past decade, a type of immunotherapy known as immune checkpoint inhibition (ICI) has been approved to treat an increasing number of cancer types. However, in spite of the profound success of these treatments in a subset of patients, most patients do not exhibit durable response following ICI therapy [1, 2]. Furthermore, the subset of patients for which these therapies are effective cannot be reliably identified prior to treatment, and the mechanisms by which some patients exhibit resistance to ICI have yet to be fully characterized [3]. An urgent need remains for biomarkers to effectively stratify patients into those likely to respond to ICI therapies and those who are not [4]. Moreover, obtaining a better understanding of mechanisms of treatment response and failure is a critical step in the development of new treatments or combinations that overcome the limitations of those currently available.

Recent research has highlighted the importance of various cell states exhibited by CD8 T cells and revealed the importance of a phenomenon known as T cell exhaustion in ICI therapy [5]. The process of CD8 T cell exhaustion, by which CD8 T cells become dysfunctional in response to prolonged antigen exposure, can be observed in the tumor immune microenvironment and in chronic viral infection and has been implicated in immunotherapy resistance [5, 6]. Since immune checkpoint inhibitors act on the inhibitory receptors that are often expressed by exhausted T cells, and successful ICI response relies on the effector function of these cells, understanding the process of T cell exhaustion is central to understanding the mechanisms of ICI response and resistance [5].

Research in mice has identified two major cell states that fall into the category of T cell exhaustion: progenitor exhausted T cells and terminally exhausted T cells [5]. Progenitor exhausted T cells are somewhat similar to memory T cells in that they are long-lived and exhibit the capacity for self-renewal. They express high levels of the protein TCF7 and low levels of the protein TOX. Terminally exhausted T cells, on the other hand, express low levels of TCF7 and high levels of TOX [5, 6]. Most of the early work to characterize the process of T cell exhaustion has been performed in mouse models, using chronic lymphocytic choriomeningitis (LCMV) infection [7, 8]. While this early work has been useful, there is a pressing need to characterize the cell states, epigenetics, and expression patterns associated with T cell exhaustion with data from human tumors rather than assuming that findings from the mouse LCMV research will generalize well to this new biological context [5]. Newer research has explored this process further in human tissue, but deeper knowledge of this process is integral to overcoming ICI therapy resistance.

Cell state changes like T cell exhaustion occur within a complex molecular context involving changes within and external to the cell. Gene-regulatory networks (GRNs) consisting of transcription factors that regulate the expression of other genes control the states and activities of cells. These, in turn, are influenced by signal transduction involving molecular cascades that may originate in other cell types. Looking at these GRNs and cell-cell signaling patterns can allow us to take a system-level view of the cell state change, rather than considering each gene in isolation.

While a standard single-cell RNA sequencing (scRNA-seq) analysis pipelines allows for the analysis of cell-intrinsic gene expression and regulation changes associated with a given cell state change, contributions of other cell types to a process within a single cell type, such as the CD8 T cell exhaustion, are challenging to infer. To address this, we developed a novel computational framework. In brief, we first inferred a pseudotemporal ordering of CD8 T cells that started with the progenitor exhausted CD8 T cells (TCF7 high, TOX low) and progressed gradually to the terminally exhausted CD8 T cells (TCF7 low, TOX high). We then calculated a sample-level CD8 T cell “exhaustion score,” based on the distribution of a sample’s CD8 T cells along this pseudotime trajectory. Subsequently, we used this score to rank samples from least to most exhausted. Cells from samples placed at the two extremes of this ranking were compared, one cell type at a time, to identify TFs that are significantly associated with the T cell exhaustion process. We applied our computational framework to scRNA-seq data generated from human melanoma samples [9]. To validate the results and identify common patterns, we also applied this framework to a basal cell carcinoma (BCC) dataset and a chronic HIV infection dataset [10, 11]. Overlap analysis between these datasets revealed common transcription factor (TF) patterns in multiple cell types in the tumor immune microenvironment associated with CD8 T cell exhaustion.

## 2 Results

### 2.1 Tumor immune microenvironment and chronic viral infection datasets

In order to investigate the relationship between CD8 T cell exhaustion and molecular activity in neighboring cell types in the tumor immune microenvironment and chronic viral infection, three publicly available single-cell RNA-seq (scRNA-seq) datasets were obtained (Tables 1, S1). The first dataset contains human melanoma tumor samples from sixteen patients [9]. The second dataset contains human BCC tumors from fourteen patients [10]. The third dataset contains peripheral blood mononuclear cell (PBMC) samples from six patients with chronic HIV infections [11]. These data were pre-processed using a standard scRNA-seq analysis pipeline (see Methods) and then analyzed with our novel pipeline, described below.

**Table 1:** scRNA-seq datasets. Three publicly available scRNA-seq datasets used – two from tumor immune microenvironment (melanoma, basal cell carcinoma) and one from chronic viral infection (HIV).

| Dataset | GEO ID | Disease | Total cells | CD8 T cells |
| --- | --- | --- | --- | --- |
| *Li et al., 2019* | GSE123814 | melanoma | 37561 | 19741 |
| *Yost et al., 2019* | GSE123139 | basal cell carcinoma | 20954 | 8136 |
| *Wang et al., 2020* | GSE157829 | HIV | 21941 | 15202 |

### 2.2 A novel computational framework to identify TFs that are significantly correlated with the T cell exhaustion process

To identify molecular mechanisms underlying the CD8 T cell exhaustion process in the tumor immune microenvironment, we have developed a novel analysis framework (Figure 1) to analyze scRNA-seq datasets. This framework consists of the following steps, applied after standard scRNA-seq pre-processing: subset to CD8+ cells, infer exhaustion pseudotime, calculate sample level exhaustion scores, infer TF activities for individual cells, conduct differential activity analysis for samples at the two extremes of the inferred T cell exhaustion trajectory, construct GRN, and then compare the patterns between different datasets. Some of the major steps in the pipeline are detailed below.

**Figure 1:**
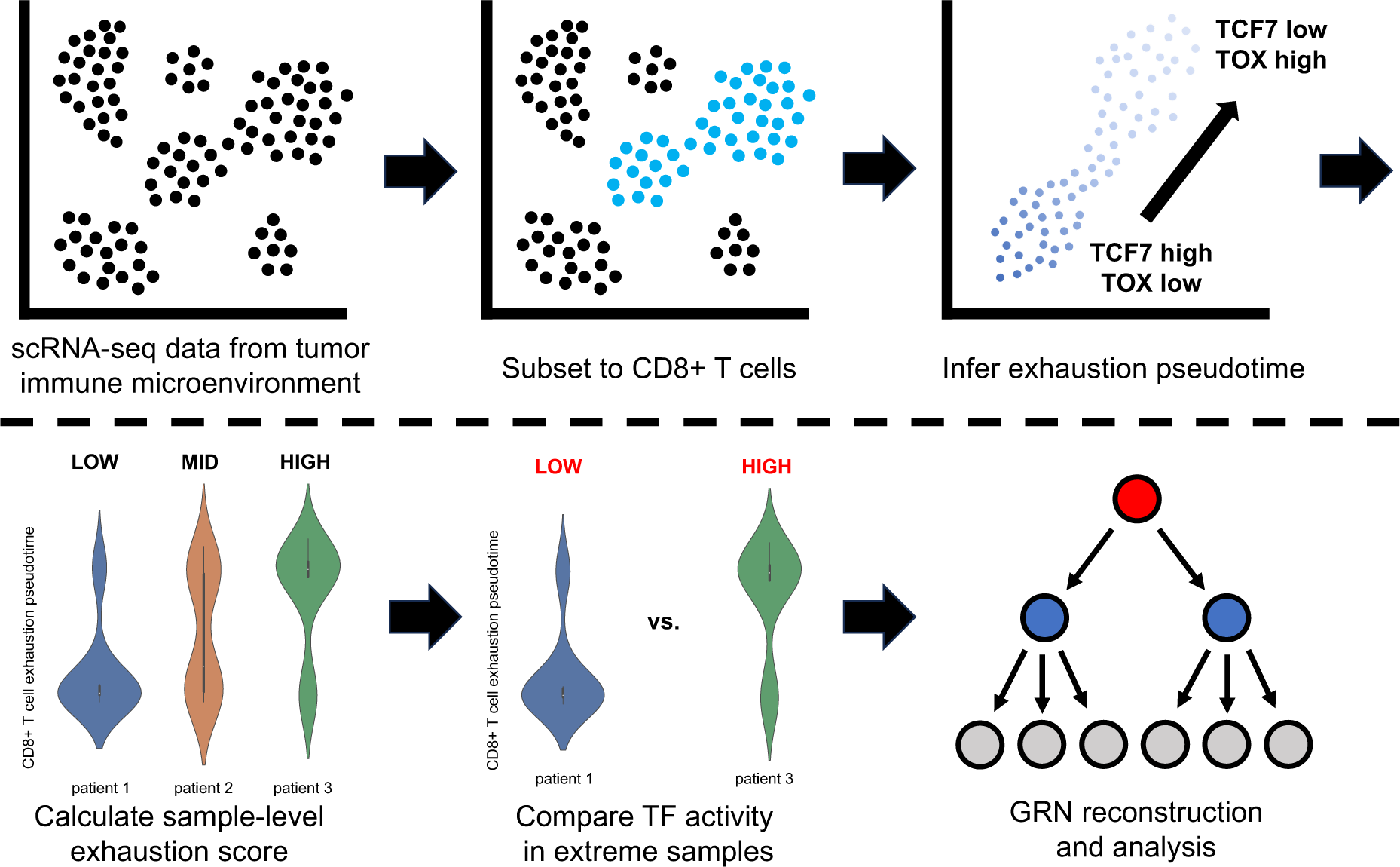
Analysis overview. A) input – scRNA-seq data from skin tumor microenvironment; B) select CD8 T cells using marker genes; C) infer pseudotime trajectory from progenitor (TCF7 high, TOX low) exhausted to terminally (TCF7 low, TOX high) exhausted CD8 T cells using Monocle3; D) calculate sample-level exhaustion score, quantifying the distribution of cells along the exhaustion trajectory by sample; E) select highest- and lowest-scoring sample groups; compare transcription factor activity in these samples for individual cell types; F) perform gene-regulatory network analysis of exhaustion-associated transcription factors.

#### 2.2.1 Inference of CD8 T cell exhaustion pseudotime trajectory

After pre-processing and preliminary analysis, the datasets were subsetted to just CD8 T cells (Figure 1B). *Monocle3* trajectory inference was then used to find a pseudotemporal ordering of cells starting from the progenitor exhausted population, which exhibited high expression of TCF7 and low expression of TOX, and proceeding to a terminally differentiated exhausted population, which exhibited low expression of TCF7 and high expression of TOX, as well as increased expression of immune checkpoint genes PDCD1, LAG3, and TIGIT (Figure 1C) [12].

#### 2.2.2 Ranking samples by sample-level CD8 T cell exhaustion score

With a cell-level CD8 T cell exhaustion trajectory, differentially expressed genes and differentially active TFs in CD8 T cells across this trajectory can be determined. However, we sought to identify molecular activity in other cell types in the tumor immune microenvironment that is significantly related to CD8 T cell exhaustion as well. To achieve this, we developed a new approach analogous to Gene Set Enrichment Analysis [13], calculating the degree of CD8 T cell exhaustion for individual patient samples according to the distribution of each sample’s CD8 T cells along this exhaustion trajectory (Figure 1D). Afterward, samples were clustered by their exhaustion score and the highest- and lowest-scoring sample groups, with CD8 T cell distributions skewed most heavily toward either the progenitor exhausted or terminally exhausted populations, were selected for downstream comparison (Figure 1E).

#### 2.2.3 Differential TF activity analysis for high- and low-exhaustion sample groups in immune cell types

Once the samples on the far ends of the CD8 T cell exhaustion spectrum were identified, TF activity in other immune cell populations was then compared between these two groups. For the melanoma dataset, TF activity in CD8 T cells, macrophages, natural killer (NK) cells, B cells, and plasma cells was considered. For the BCC dataset, TF activity in CD8 T cells, macrophages, and NK cells was considered. For the HIV dataset, we looked at CD8 T cells, macrophages, B cells, and plasma cells.

### 2.3 Application of the computational framework to scRNA-seq datasets from human skin cancer and HIV infection

We first applied this novel computational framework to a human melanoma scRNA-seq dataset (Figure 2) [9]. Once the CD8 T cells were identified, a pseudotime trajectory was inferred (Figure 2C, S2-4). The CD8 T cells in this dataset have a TCF7-high state (Figure 2A), where pseudotime starts, and transition gradually to a TOX-high state (Figure 2B). This transition also shows a progressive increase in expression of immune checkpoint genes PDCD1, LAG3, and TIGIT (Figures 2D-F, S3). The sample-level T cell exhaustion score analysis showed that samples p23, p25, and p27 have the highest T cell exhaustion scores and samples p1 and p24 have the lowest exhaustion scores (Figure 2G). The differential TF activity between these two sample groups (as quantified by *pyscenic* regulons and *AUCell*) was then compared.

**Figure 2:**
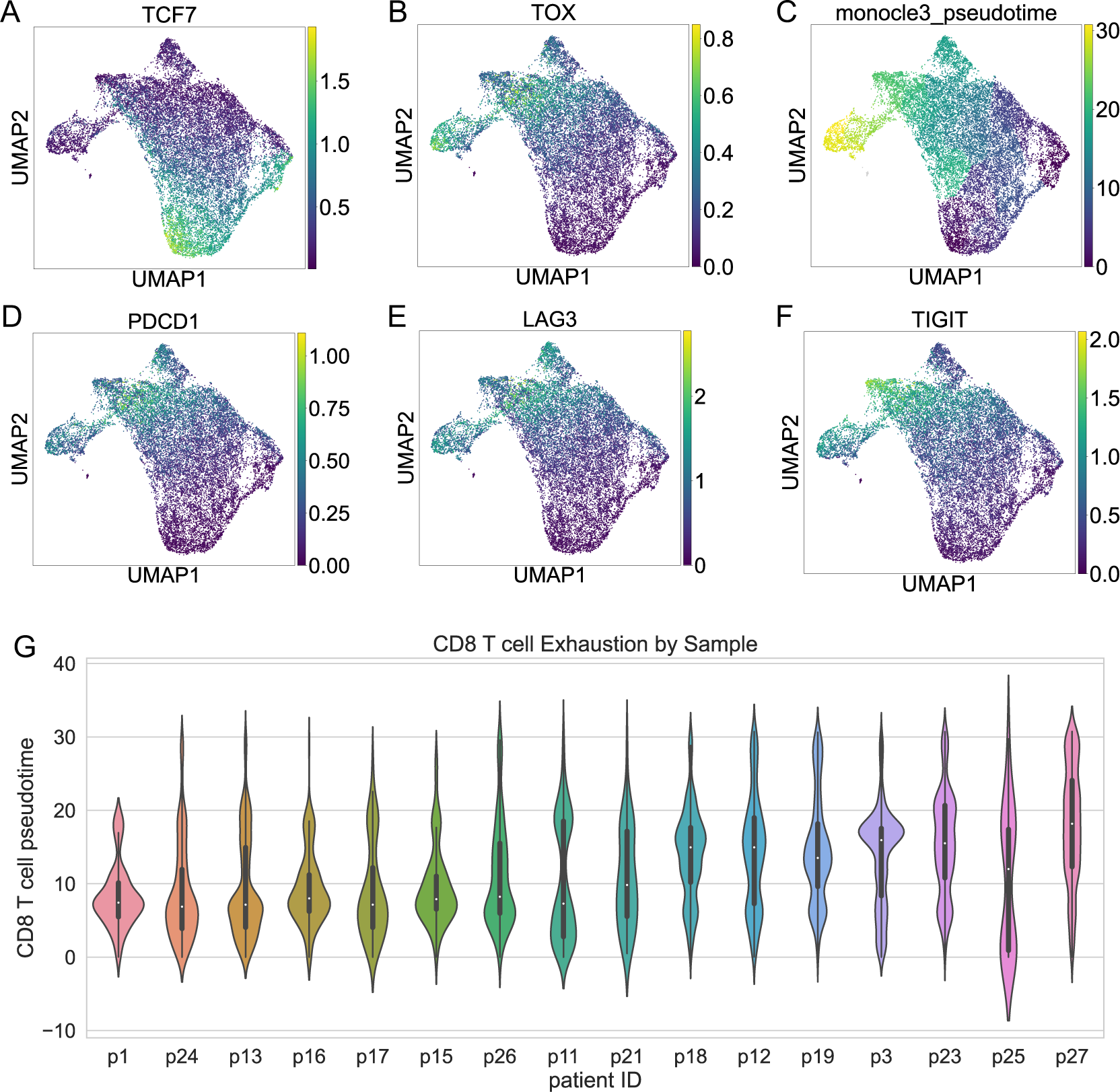
CD8 T cell exhaustion in human skin tumor samples. A, B, D-F) imputed gene expression of progenitor exhausted marker TCF7, terminal exhaustion marker TOX, and immune checkpoints LAG3, TIGIT, and PDCD1; C) Monocle3 pseudotime, characterizing progression from progenitor exhausted to terminally exhausted CD8 T cells; G) exhaustion pseudotime of CD8 T cells, ordered by sample-level exhaustion score.

We also applied this computational framework to a BCC dataset to investigate the generality of this pattern in the skin tumor microenvironment. Furthermore, we applied the framework to a chronic HIV infection dataset as well (Figures S1, 3). Chronic viral infection data from human patients was included to enable the comparison of the T cell exhaustion process between the human tumor and viral infection contexts. The similarities and differences in the etiology and regulation of this cell state change between these two biological contexts is not yet fully clear. To address this, we compare and contrast the molecular profiles between the two.

A similar trajectory of CD8 T cell state change was observed in these two datasets (Figures S1C, 3C, S5-10). We observed a similar trajectory of key markers of stemness, differentiation, and immune checkpoints (Figure S6, S9). For the BCC data, fourteen patient samples were scored and ranked, the high exhaustion group (samples su008, su013, su014) was compared against the low exhaustion group (002, 007) and differential TF activity between these groups was considered in each cell type (Figure S1G). For the chronic HIV data, six patient samples were scored and ranked and the high (samples 4 and 5) and low (samples 2 and 3) exhaustion groups were similarly compared (Figure 3G).

**Figure 3:**
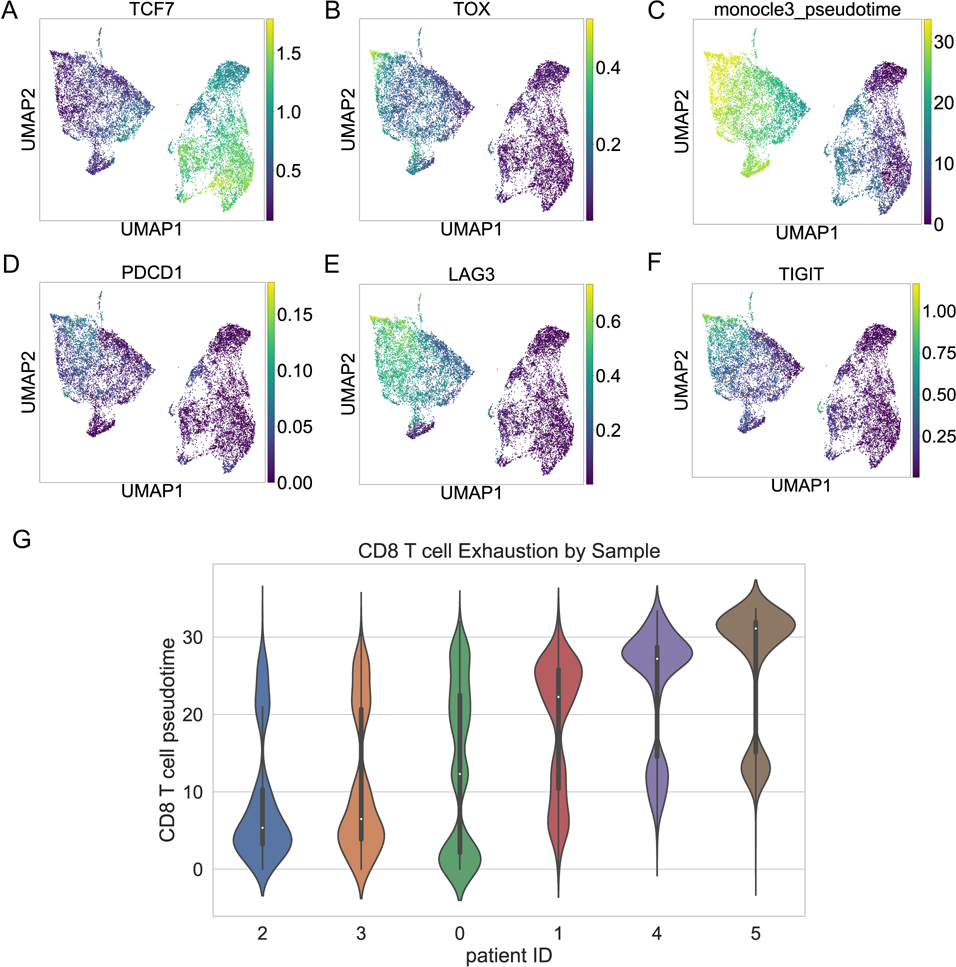
CD8 T cell exhaustion in human HIV samples A, B, D-F) imputed gene expression of progenitor exhausted marker TCF7, terminal exhaustion marker TOX, and immune checkpoints LAG3, TIGIT, and PDCD1 C) Monocle3 pseudotime, characterizing progression from progenitor exhausted to terminally exhausted CD8 T cells G) exhaustion pseudotime of CD8 T cells, ordered by sample-level exhaustion score.

### 2.4 Identification of gene regulatory patterns shared across skin tumor datasets that are related to T cell exhaustion

To uncover the cell type-specific transcription factor (TF) activity associated with the CD8 T cell exhaustion process that is consistently found in the skin tumor microenvironment, overlap analysis was performed via Fisher’s exact test for results obtained from two skin cancer datasets (Methods). Cell types shared across both skin cancer datasets, including CD8 T cells, macrophages, and NK cells, were considered in the overlap analysis. We found significant overlap in TFs up-regulated (44 shared, adjusted p-value = 1.24×10^-11^) and down-regulated (68 shared, adjusted p-value = 5.42×10^-13^) in the CD8 T cells of the most exhausted samples, suggesting a shared mechanism in the T cell exhaustion process across skin tumor microenvironments (Figure 4A, B). Additionally, there was significant overlap in TFs up-regulated (25 shared, adjusted p-value = 1.69×10^-6^) in the macrophages of the most exhausted samples, as well as up- (23 shared, adjusted p-value = 8.59×10^-12^) and down-regulated (27 shared, adjusted p-value = 5.10×10^-5^) TFs in the NK cells (Figure 4C-F). However, we failed to observe a significant overlap of TFs down-regulated in macrophages from the most exhausted skin tumor microenvironments, which may suggest that the exhaustion-associated activity in macrophages mostly involves signaling pushing CD8 T cells toward the terminally exhausted state rather than holding them in the progenitor state (Figure 4D).

**Figure 4:**
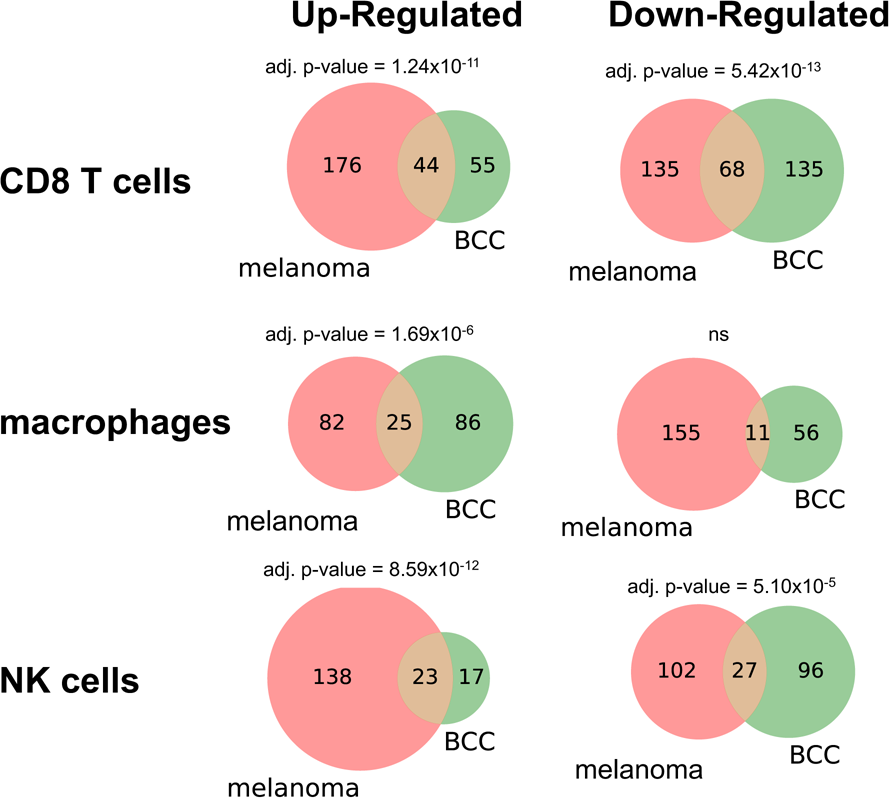
Overlap in exhaustion-associated transcription factor activity between melanoma and basal cell carcinoma (BCC) datasets Overlap analysis was performed between melanoma and basal cell carcinoma datasets to identify the proportion of significantly exhaustion-associated TFs that were shared vs. non-shared and whether this overlap was significant. A-C, E-F) The two tumor datasets exhibit significant overlap in up- and down-regulation of CD8 T cell activity, up-regulation of macrophage activity, and up- and down-regulation of NK cell activity associated with CD8 T cell exhaustion. D) The overlap of TFs down-regulated in the most exhausted samples was not significant.

#### 2.4.1 Transcription factors up-regulated in terminally exhausted tumor-infiltrating CD8 T cells

We observed the up-regulation of certain CD8 T cell-specific transcription factor activity in the most exhausted samples in both tumor datasets, including IRF2, CTCF, E2F1, E2F2, E2F8, ETV7, and EOMES. Some of these have been found to be related to the CD8 T cell exhaustion process in previous experiments. IRF2 expression in tumor-infiltrating CD8 T cells has been shown to drive T cell exhaustion [14]. It pushes CD8 T cells toward the terminal exhaustion state by acting on TOX [15]. CTCF works with TCF1 as a cofactor to promote proliferative homeostasis in CD8 T cells [16]. It has been found to control the relative abundance of terminally exhausted CD8 T cells [17].

Members of the E2F transcription factor family are known cell cycle regulators. E2F1 is a regulator of cell cycle and proliferation; however, it may not be necessary for progression to the exhausted CD8 T cell state, based on viral experiments in mice [18]. E2F2 and E2F8 expression has been found in exhausted T cells in COVID-19 infection [19]. Additionally, E2F2 is most active in PD1+/CD39+ CD8 T cells [20].

ETV7 drives cell proliferation in some cell types [21, 22]. Some previous findings indicate that chronically stimulated CD8 T cells show ETV7 activity that down-regulates inflammation, potentially via ETS1 and TBX21, which we also found to be significantly up-regulated in the CD8 T cells of the most exhausted tumor samples in both datasets [23]. EOMES, which we found to be up-regulated in the most exhausted tumor samples, is a key regulator of CD8 T cell exhaustion as well, and binds competitively to the same sites as T-Bet (TBX21); however, the exact roles and functions of these factors is not yet clear [24, 25, 26, 27]. ETS1 is known to be important in regulating T cell state [28, 29]. Additionally, ETS1 knockdown leads to increased anti-tumor cytotoxic activity of CD8 T cells [30].

#### 2.4.2 Transcription factors down-regulated in terminally exhausted tumor-infiltrating CD8 T cells

We also observed the down-regulation of certain CD8 T cell-specific transcription factor activity in the most exhausted samples in both tumor datasets, including LYL1, BCL11A, PLAG1, and BCL-6. Several of these are known to be responsible for maintaining the stem-like state of the progenitor exhausted CD8 T cells. LYL1 is responsible for stem / progenitor state and self-renewal in thymocytes and T cells [31, 32, 33]. BCL11A slows differentiation of T cells to maintain self-renewal [34]. It is required for normal lineage development of lymphocytes [35, 36]. PLAG1 helps to maintain self-renewal in hematopoietic stem cells [37]. BCL-6, acting in opposition to PRDM1 (encodes BLIMP-1), maintains the relative proportion of progenitor (as opposed to terminally differentiated) exhausted CD8 T cell populations; TCF1, which drives self-renewal in progenitor exhausted CD8 T cells, activates BCL-6 and suppresses PRDM1 [38, 39].

#### 2.4.3 Down-regulation of Kruppel-like factor (KLF) signaling in terminally exhausted tumor-infiltrating CD8 T cells

We observed down-regulation of KLF2, KLF3, and KLF4 signaling in the CD8 T cells of the most exhausted melanoma samples, with KLF3 and KLF4 down-regulated in the BCC samples as well. This activity concords with the findings of previous mouse LCMV research [40]. The most exhausted HIV samples also exhibited down-regulation of KLF3 and KLF4; however, they showed up-regulation of KLF2, as did the BCC data.

It is worth noting that KLF4 has been linked to increased effector function in these terminally exhausted CD8 T cell subsets [41]. This concords with the finding that the most exhausted samples exhibited down-regulation of KLF4 in their CD8 T cells and also had poorer response to immunotherapy. We also observed down-regulation of AP-1 signaling factors such as FOS, FOSB, JUN, JUNB in CD8 T cells across multiple datasets, the activity of which has been linked to KLF4 signaling in other work [41].

### 2.5 Transcription factors up-regulated in macrophages of highly exhausted tumor samples

In addition to identifying TF activity in CD8 T cells associated with T cell exhaustion, some of which was in concordance with previous findings, we also identified TF activity in macrophages associated with the degree of CD8 T cell exhaustion in a sample. In both tumor datasets, significant up-regulation of TBX21 (T-Bet), PRDM1, RUNX3, CREM, IRF1, FOXP3, STAT2, NFKB1 and NFKB2 was observed. Some of these transcription factors, including TBX21 and PRDM1, also exhibit up-regulation in the most exhausted CD8 T cells themselves, suggesting that these two cell types may be responding to some of the same signals in the exhausted tumor immune microenvironment. Significant up-regulation of NFKB subunits NFKB1 and NFKB2 was observed in the macrophages of the most exhausted samples across all three datasets. Additionally, NFKB subunit RELB was up-regulated in the most exhausted BCC and HIV samples, with RELA up-regulated in the melanoma macrophages. These results suggest the up-regulation of NF Kappa-B signaling in macrophages of both highly exhausted tumor and HIV patients.

### 2.6 Gene-regulatory network modules associated with CD8 T cell exhaustion

#### 2.6.1 Exhaustion-associated CD8 T cell GRNs

To better understand the regulatory relationships among TFs associated with CD8 T cell exhaustion, we reconstructed gene-regulatory network modules using a subset of the highly exhaustion-associated TFs whose activity in CD8 T cells was up-regulated in the most exhausted samples (Figure 5A). Regulatory relationships between these significant TFs were inferred using *pyscenic* gene regulatory network inference, which is based on co-expression between a transcription factor and its potential target followed by search for a TF’s binding motif in the promoter region of a potential target (Methods). Edge widths represent the strength of the inferred relationship. We identified a larger module consisting primarily of tumor-specific exhaustion-associated TF regulation, which include TBX21 and EOMES. We also identified two up-regulated GRN modules comprising viral-specific exhaustion-associated TFs. In one, EGR1 activates JUN (forms AP-1 complex with FOS), which then activates ATF5, BPTF, and KLF6. The other includes BCL11A, PAX5, POU2F2, and SOX5.

**Figure 5:**
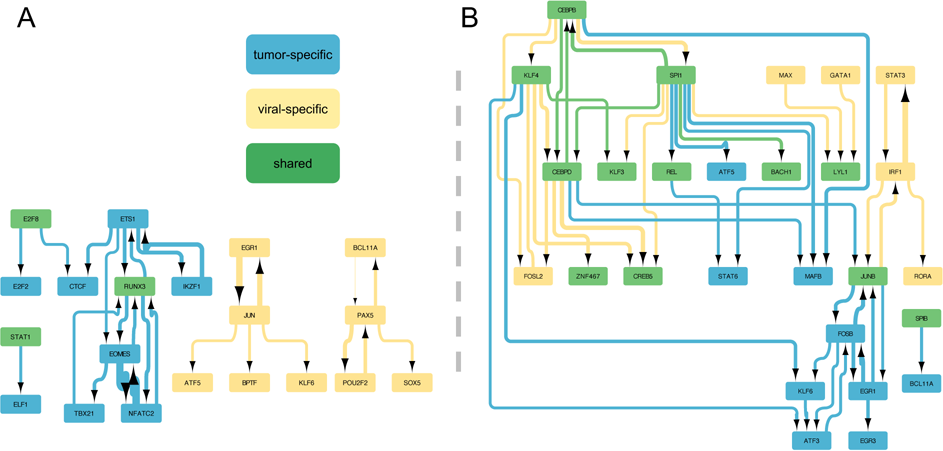
Exhaustion-related transcription factor activity in CD8 T cells Exhaustion-associated transcription factors (graph nodes) and their regulatory relationships, inferred with pyscenic (graph edges) A) transcription factors up-regulated in the CD8 T cells of the most CD8 T cell-exhausted immune microenvironments B) transcription factors down-regulated in the CD8 T cells of the most CD8 T cell-exhausted immune microenvironments.

The same GRN module reconstruction was performed for TFs whose activity was found to be down-regulated in the CD8 T cells of the most exhausted samples (Figure 5B). This module included some TF down-regulation shared across the tumor and viral contexts, including KLF3 and KLF4, as well as tumor- and viral-specific activity. Interestingly, KLF6 was up-regulated in the most exhausted viral samples (adjusted p-value: 0.0401) but down-regulated in the most exhausted tumor samples (adjusted p-values for melanoma, BCC: 7.34×10^-7^, 6.51×10^-54^), suggesting a potential difference in mechanism behind these cell state trajectories.

#### 2.6.2 Exhaustion-associated macrophage GRNs

GRN module reconstruction was also performed for exhaustion-associated TF activity in macrophages (Figure 6). We identified a large GRN module of up-regulated TF activity in viral patients, including TFs related to FOS/JUN/AP-1 signaling, which was connected to an NFKB module via IRF1. This NFKB / IRF1 activity was found to be up-regulated in the most exhausted samples across both tumor datasets and the viral dataset, indicating that NFKB activity in macrophages may be associated with CD8 T cell exhaustion in both disease contexts. We also observed tumor-specific up-regulation of TBX21 and RUNX3, whose activity is associated with exhaustion in the CD8 T cells themselves as well. A number of transcription factors and complexes were found to be significantly up-regulated in the macrophages of the most exhausted samples across all 3 datasets, including NFKB, IRF1, BHLHE40, FOSL2, NFIL3, and CREM. Previous research indicates that NFKB regulates IRF1 [42, 43]. Additionally, there is cross-regulation among NFKB, AP-1, and IRF1 signaling [43]. We also identified two viral-specific GRN modules of TFs down-regulated in the macrophages of the most exhausted samples, which included members of the interferon regulatory factor (IRF2, IRF5, IRF9) and STAT (STAT1, STAT2) families.

**Figure 6:**
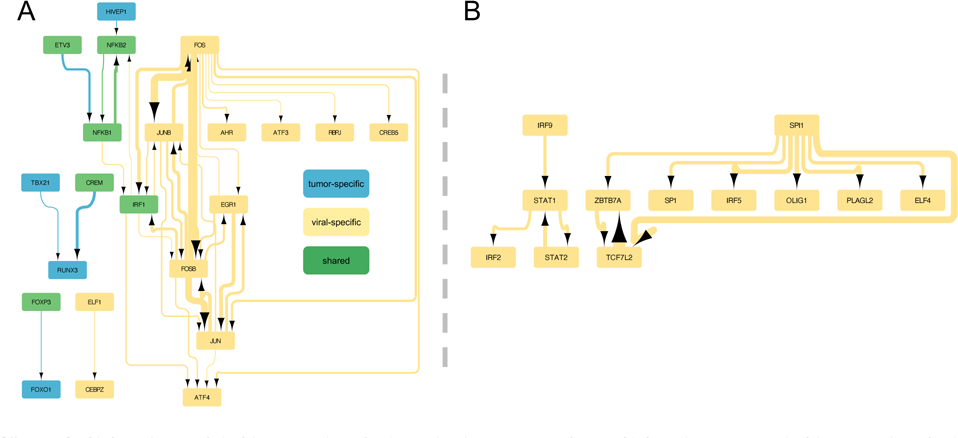
Exhaustion-related transcription factor activity in macrophages Exhaustion-associated transcription factor activity in macrophages (graph nodes) and their regulatory relationships, inferred with pyscenic (graph edges) A) transcription factors up-regulated in the macrophages of the most CD8 T cell-exhausted immune microenvironments B) transcription factors down-regulated in the macrophages of the most CD8 T cell-exhausted immune microenvironments.

### 2.7 Identification of pathways significantly related to T cell exhaustion in macrophages

In order to identify biological pathway activity in macrophages that may play a significant role in the exhaustion process, we performed pathway enrichment analysis on differentially expressed genes in the macrophages of high- and low-exhaustion samples using *Reactome* (Figure 7, Methods) [44, 45, 46, 47]. We observed some tumor-specific exhaustion-associated pathways in macrophages, including a number of immune pathways. We saw an increase in TNFR2 non-canonical NF-kB signaling in the tumor context and an increase in p75NTR signals via NF-kB in the viral context, which aligns with our observation of an up-regulation of NFKB1 and NFKB2 in the most exhausted samples across the tumor and viral contexts, but indicates that the mechanism here may be slightly different.

**Figure 7:**
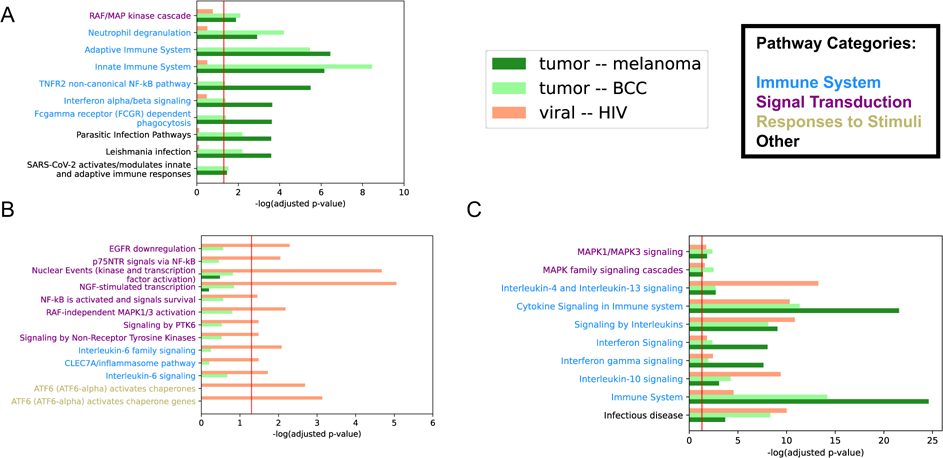
Pathway analysis results of exhaustion-associated DEGs in macrophages. Pathways enriched in genes over-expressed in macrophages of high-exhaustion samples relative to low-exhaustion samples; red line = adjusted p-value cutoff of 0.05 A) tumor-specific exhaustion-associated pathway activity B) viral-specific exhaustion-associated pathway activity C) exhaustion-associated pathway activity shared between tumor and viral contexts.

In both tumor and viral infection, we observed up-regulation of the interleukin 10 signaling pathway (Figure 7C). Importantly, we see up-regulation of NFIL3 in the macrophages of high-exhaustion samples (see above). Published results show that NFIL3 drives IL-10 activity, further supporting the possible causal relationship between NFIL3 and the IL-10 pathway [48, 49]. While the role of IL-10 on CD8 T cell state and activity appears to be multifaceted, this anti-inflammatory cytokine may drive exhaustion of CD8 T cells via signaling by PRDM1, which was identified as an up-regulated TF in the high-exhaustion macrophages [50, 51, 52]. We also observed some MAPK-related signaling pathways up-regulated in exhaustion-associated macrophages, such as MAPK family signaling cascades, RAF/MAP kinase cascades, MAPK1/MAPK3 signaling, and RAF-independent MAPK1/3 activation. In addition to this, we also observed up- and down-regulation of a number of immune, cell cycle-related, and other pathways in the CD8 T cells themselves (Figures S14-19).

## 3 Discussion

The promising yet incomplete success of immune checkpoint inhibitors has led to significant interest in and scrutiny of the CD8 T cell exhaustion process due to its association with treatment outcome. Progress in this research area can change the way cancer and chronic infection are treated. This process has garnered much attention in recent years as many attempts to find ways to modulate this process, with the hope of improving immunotherapy outcomes and expanding the population of potential responders to this class of treatments through the development of drugs that modulate the tumor immune microenvironment to improve the likelihood of response to ICI therapy.

To better understand the molecular mechanisms of the CD8 T cell exhaustion process, we have developed a new computational approach to analyze scRNA-seq datasets collected from tumor immune microenvironments. The advantage of this framework lies in its ability to consider the potential contribution of gene or TF activity in multiple cell types to a change in cell state in a cell type of interest, rather than only considering activity within that single cell type. Existing tools allow us to determine which genes and transcription factors are differentially active across the exhaustion pseudotime trajectory; our approach produces a sample-level exhaustion score, therefore allowing the differential activity analysis of transcription factors in other cell types along with CD8 T cells. By applying our novel computational framework to single-cell transcriptomics data from human tumor and viral infection samples, we identified the potential contribution of various immune cell types to this process. This work contributes to ongoing efforts that seek to understand how, when, and why ICI therapy succeeds and fails and, ultimately, to improve clinical outcomes in patients with late-stage cancers.

Our novel computational approach successfully identified TFs in CD8 T cells associated with CD8 T cell exhaustion, implying their potential involvement in immune checkpoint inhibitor response. Many of these TFs recapitulate published results from previous investigations of this process. By examining the literature, we verified the importance of transcription factors such as TBX21 (T-bet), BLIMP1, the E2F family, and the KLF family. More importantly, this approach also identified TFs in other cell types, including macrophages, which sheds light on the state of the surrounding tumor immune microenvironment and how it may contribute to CD8 T cell exhaustion. In the macrophages of samples with the greatest degree of CD8 T cell exhaustion, across both tumor datasets and the HIV data, we found up-regulation of NFKB signaling, as well as IRF1, BHLHE40, FOSL2, NFIL3, and CREM. Some of these TFs form a gene-regulatory network module (Figure 6A). Additionally, NFIL3 has the potential to regulate the activity of a key cytokine, IL-10, which has been linked to the CD8 T cell exhaustion process by previous research [50, 51, 52]. The activity of cytokines like IL-10 may partially explain the association between the abundance of macrophages in the tumor immune microenvironment and ICI response [53]. In addition to the up-regulation of IL-10 regulator NFIL3 itself in exhaustion-associated macrophages, we also observed up-regulation of genes in the IL-10 signaling pathway in exhaustion-associated macrophages. This provides further support for the association between IL-10 signaling in the tumor immune microenvironment and CD8 T cell exhaustion. The activity of this anti-inflammatory cytokine as it relates to CD8 T cell exhaustion is worthy of further investigation. By identifying cell type-specific TF activity in the tumor immune microenvironment associated with the degree of CD8 T cell exhaustion in a patient sample, and by organizing these relevant TFs into a gene-regulatory network that can be studied at the system level, we contribute to a clearer understanding of this critical biological process.

This work fulfills the need to compare and contrast the CD8 T cell exhaustion process between the tumor immune microenvironment and the chronic viral infection context in human tissue as well. We found some exhaustion-associated GRN modules in CD8 T cells and macrophages that were viral-specific, while others were a combination of tumor-specific and shared. As these results come from human data, they are more relevant to the human disease context than earlier findings from LCMV in mice, which they may confirm or contradict. In CD8 T cells, we observed up-regulation of two viral-specific GRN modules, with one related to AP-1 signaling. We also observed viral-specific up-regulation of AP-1-related TF activity in exhaustion-associated macrophages, as well as down-regulation of an IRF9-regulated GRN module. One caveat to bear in mind when comparing these tumor and viral samples is that the difference in tissue context (peripheral blood vs. skin tumor microenvironment) likely drives some of the observed differences in expression patterns.

In addition to providing insights into the CD8 T cell exhaustion process, this computational pipeline can also be applied to other biological questions. This approach allows researchers to consider the potential contribution of other cell types to a cell state change of interest, whether in cancer, development, or another biological context. While we used this functionality to identify potential drivers of CD8 T cell exhaustion throughout the tumor immune microenvironment, it can be applied to any process wherein a continuous trajectory of cell states can be identified in scRNA-seq data and mapped in pseudotime. By moving to the sample level and ranking samples based on the distribution of their cells across a trajectory, we build on currently available methods to add this important perspective.

To summarize, this work illuminates some key differences between T cell exhaustion in human tumors and chronic infections and reveals gene-regulatory networks in immune cell populations within a tumor that may mediate response to immune checkpoint inhibition therapy. The novel framework we developed also allows researchers to consider the potential contribution of other cell types to a cell state change of interest, giving it wider applicability in the field of single-cell transcriptomics research.

## 4 Data Availability

The three scRNA-seq datasets used in this paper have been previously published, and are available via Gene Expression Omnibus: *GSE123814* [9], *GSE123139* [10], *GSE157829* [11] – https://www.ncbi.nlm.nih.gov/geo/

## 5 Code Availability

Code used in this paper can be located in the following GitHub repository:

https://github.com/christopher-klocke/exhaustion-microenvironment

## 6 Results Availability

Hosted at *zenodo* with DOI: https://doi.org/10.5281/zenodo.10088918

## Acknowledgements

This research was supported by the US NLM Training Grant (*T15-LM007088*, CK) and US NIH (*U41 HG003751*, *U24HG012198* and *U01CA239069*, GW) grants. The authors would also like to thank Brett Davis and Nathaniel Evans for discussion of original ideas.

## 7 Methods

### 7.1 Datasets

Publicly available datasets were downloaded from Gene Expression Omnibus using the following identifiers: *GSE123814* [9], *GSE123139* [10], *GSE157829* [11]. The *Li et al.* dataset contains human melanoma samples from 16 patients. The *Yost et al.* dataset contains human basal cell and squamous cell carcinoma samples from 15 patients. The *Wang et al.* dataset contains human peripheral blood mononuclear cell (PBMC) samples from six HIV-infected patients. For the *Yost et al.* dataset, only pre-treatment (with immune checkpoint inhibition) samples were used.

### 7.2 Pre-processing, dimensionality reduction, batch correction, and clustering

Data files were unzipped and converted to the *AnnData* format [54] using in-house Python and Bash scripts and stored as .h5ad files. Datasets were subsetted to just protein-coding genes per *UniProt* [55]. Pre-processing was performed using the standard *scanpy* (*1.9.1*) [54] pipeline for scRNA-seq data, including quality filtering, normalization, and log-transformation steps. Doublets were identified and removed using *DoubletDetection* (*4.2*) [56]. Dimensionality reduction was performed with PCA. In order to account for potential batch effects between samples, batch correction was performed with *Batchelor* (*1.14.1*) [57]. Clustering was performed using the Leiden algorithm (*leidenalg 0.8.10*) [58]. The dimensionality was further reduced to two dimensions for visualization using UMAP (*umap-learn 0.5.3*) [59, 60].

### 7.3 Cell cluster annotation

*MAGIC* (*magic-impute 3.0.0*) [61] was used to impute expression for marker genes collected from databases and the literature. Cell types were manually assigned to clusters based on the following marker genes for the *Li et al.* and *Wang et al.* data sets: CD8 T cells: CD3, CD8; CD4 T cells: CD3, CD4; regulatory T cells: CD3, CD4, FOXP3; macrophages: CD68, CD163, CD14; natural killer cells: CD56, CD16, CD3, KLRB1, NKG7, NKG2A, GZMB, NCAM1; B cells: CD19, CD27, CD38; plasma cells: SDC1, CD20. We used the previous cell type annotation for *Yost et al.* dataset.

### 7.4 Inference of CD8 T cell exhaustion trajectory with *Monocle3*

Datasets were subsetted to just CD8 T cell clusters for trajectory inference. A pseudotemporal ordering of cells from progenitor exhausted to terminally exhausted was inferred for CD8 T cells using *Monocle3* (*1.3.1*) [12]. Root cells were chosen from the principal graph based on expression of key exhaustion-associated genes, including TCF7 (high), TOX (low), AND LAG3 (low).

### 7.5 Calculation of sample-level exhaustion scores

To assess the distribution of a sample’s CD8 T cells along the exhaustion pseudotime path relative to the overall distribution (across all samples), a cell enrichment approach was used by adopting the Gene Set Enrichment Analysis (GSEA) method (*gseapy 0.9.5*) [13]. Whereas in GSEA, a gene set is queried for enrichment at one end of a ranking of genes, we quantified the degree to which a subset of cells (derived from a given sample) was enriched at one end of a ranking of cells (exhaustion pseudotime across all samples). The resulting score quantifies the degree to which a sample’s CD8 T cells are biased toward the progenitor exhausted or terminally exhausted state.

### 7.6 Sample clustering and selection of extremes

We used the Jenks natural breaks algorithm (*jenkspy 0.3.2*) to divide the samples into clusters based on similar exhaustion scores [62]. Sensitivity analyses were performed to ensure reasonable robustness to the choice of cluster number. The sample clusters with the highest and lowest scores were chosen for further analysis. By comparing the extreme samples from the ends of the distribution, we increased power to detect changes in other cell types associated with these different immune microenvironments. Additionally, by using multiple samples from each end, we minimized the effect of individual patient background and increased the number of cells used to calculate the results. For every dataset, at least two samples were used at each end of the sample ranking.

### 7.7 Quantifying transcription factor regulon activity with *SCENIC*

We used the *SCENIC* gene-regulatory network inference algorithm, implemented in the *pyscenic* (*0.11.2*) Python library. Briefly, *GRNBoost* was used to infer correlation modules, which were then pruned by *CisTarget* to obtain regulons, each consisting of a transcription factor and its inferred, directly regulated targets (*arboreto 0.1.6*). The activity of each regulon was then quantified for each cell using *AUCell* [63, 64, 65].

### 7.8 Comparison of transcription factor activity between exhaustion-low and exhaustion-high samples in each cell type

*AUCell* scores for a given regulon were used as a proxy for the activity of the corresponding transcription factor. To determine whether a given transcription factor was significantly up- or down-regulated in a given cell type in one sample group relative to the other, the following comparison was made: using a Mann-Whitney U test (*scipy 1.9.0*), the distributions of TF activity scores were compared between the two sample groups. Moving to the sample level for this comparison allows for the assessment of potentially exhaustion-associated activity in other cell types within the tumor immune microenvironment. P-values were then corrected using the Benjamini-Hochberg method.

### 7.9 Overlap analysis

Once exhaustion-associated cell-type specific TF activity was identified for each dataset, the results were compared across datasets. We used a Fisher’s exact test to perform an overlap analysis, determining whether a given cell type had shared significant up- or down-regulated TFs between two datasets. The resulting p-values were corrected using the Bonferroni method.

### 7.10 Analysis of exhaustion-associated gene-regulatory sub-networks

TF-target relationships inferred using *pyscenic* were used to connect significant TFs into gene-regulatory sub-networks. Two thresholds were then used to generate the GRN figures. Importance scores were exported from the *GRNBoost* results for each TF-target relationship. If both tumor datasets had an inferred GRN graph edge for a TF-target relationship at a score above 1 but the viral dataset did not, the edge was labeled as tumor-specific. If the viral score was above one but neither of the tumor scores were above 1, it was listed as viral-specific. If all 3 had a score above 1, the edge was considered to be shared across these biological contexts. The GRN was then pruned using the higher threshold; if the maximum score for a potential TF-target edge exceeded the higher threshold, the edge and its associated nodes were included in the final network visualization. This threshold varied from 25 to 30 for different GRN figures. All GRN visualizations were created using *Cytoscape* (*v3.9.1*) [66]. A subset of the top-scoring TFs were displayed in the GRN figures; a full list of exhaustion-associated TFs and their p-values can be found in the Supplemental Results files.

### 7.11 Pathway enrichment analysis with *Reactome*

Using *Reactome* (*v85*) pathway gene sets and pathway hierarchy, we performed a pathway enrichment analysis on genes differentially expressed (per two-sided Mann-Whitney U test, *scipy*) in CD8 T cells and macrophages between the high- and low-exhaustion samples [44, 45, 46, 47]. A Fisher’s exact test was used to quantify overrepresentation of a pathway’s genes in these differentially expressed genes. Out of 20376 total protein-coding genes (per *UniProt v2022_01*), any gene with an adjusted p-value less than 0.05 was considered to be differentially expressed. These were then tested for overrepresentation in each pathway’s gene set.

### 7.12 Other package versions

*matplotlib=3.5.1; matplotlib-base=3.5.1; networkx=2.8.5; numpy=1.22.3; numpy-base=1.22.3; pandas=1.4.2; python=3.10.4; scikit-learn=1.1.2; seaborn=0.11.2*

