## Supplemental for "Identification of Cellular Interactions in the Tumor Immune Microenvironment Underlying CD8 T Cell Exhaustion"

### SUPPLEMENTAL MATERIALS

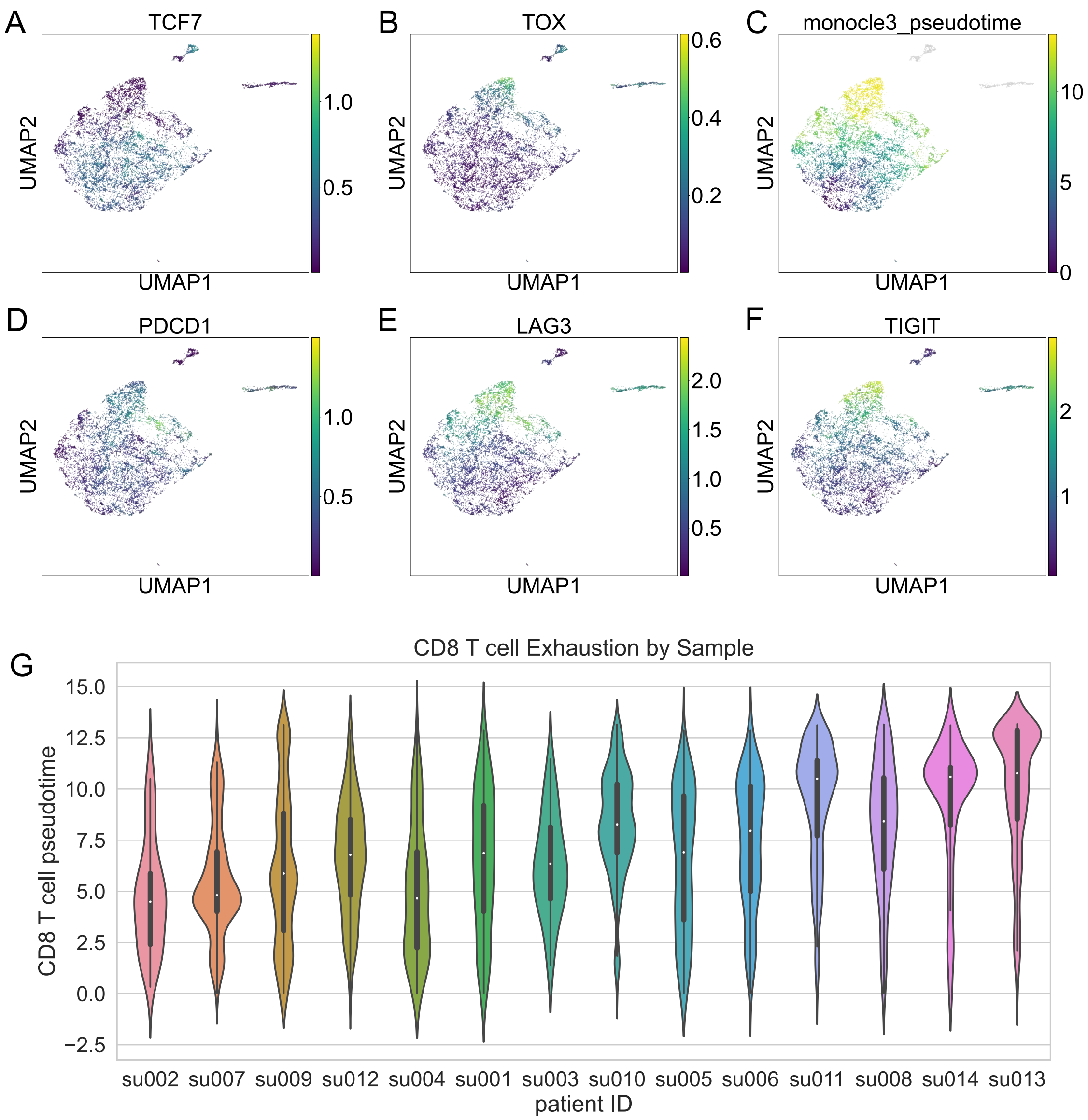


**Figure S1**: **CD8 T cell exhaustion in human skin tumor (basal cell and squamous cell carcinoma) samples**
**a, b, d-f)** imputed gene expression of progenitor exhausted marker TCF7, terminal exhaustion marker TOX, and immune checkpoints LAG3, TIGIT, and PDCD1 **c)** Monocle3 pseudotime, characterizing progression from progenitor exhausted to terminally exhausted CD8 T cells **g)** exhaustion pseudotime of CD8 T cells, ordered by sample-level exhaustion score

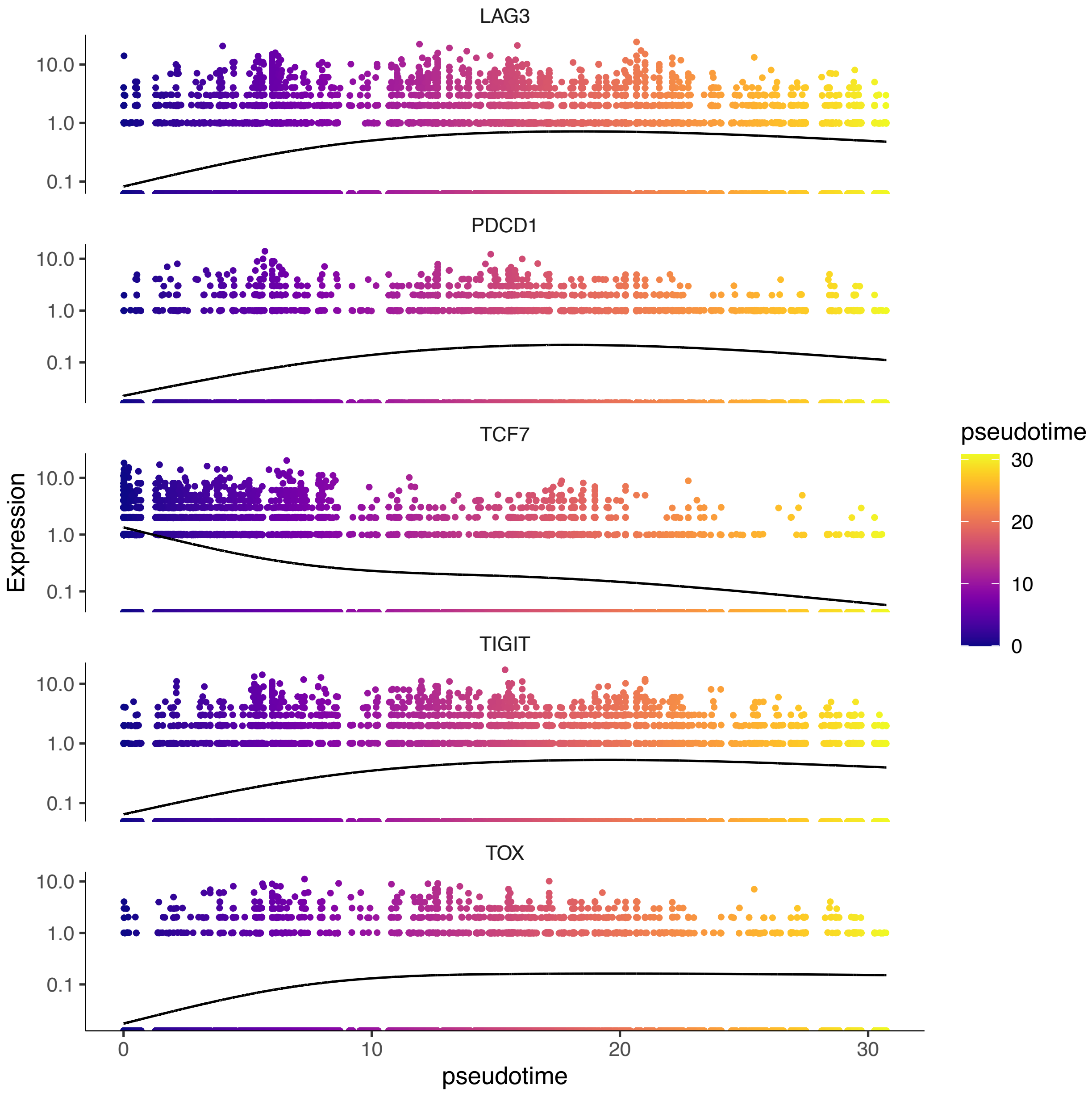


**Figure S2**: **gene expression vs. pseudotime in melanoma dataset**
Expression of TCF7, the primary marker of the progenitor exhausted CD8 T cell state, starts high and is monotonically decreasing across pseudotime. TOX, the primary marker of the terminally exhausted CD8 T cell state, shows the opposite relationship. Immune checkpoint genes LAG3, PDCD1, and TIGIT show broadly similar expression patterns to TOX relative to pseudotime, although they decrease slightly for the highest pseudotime values.

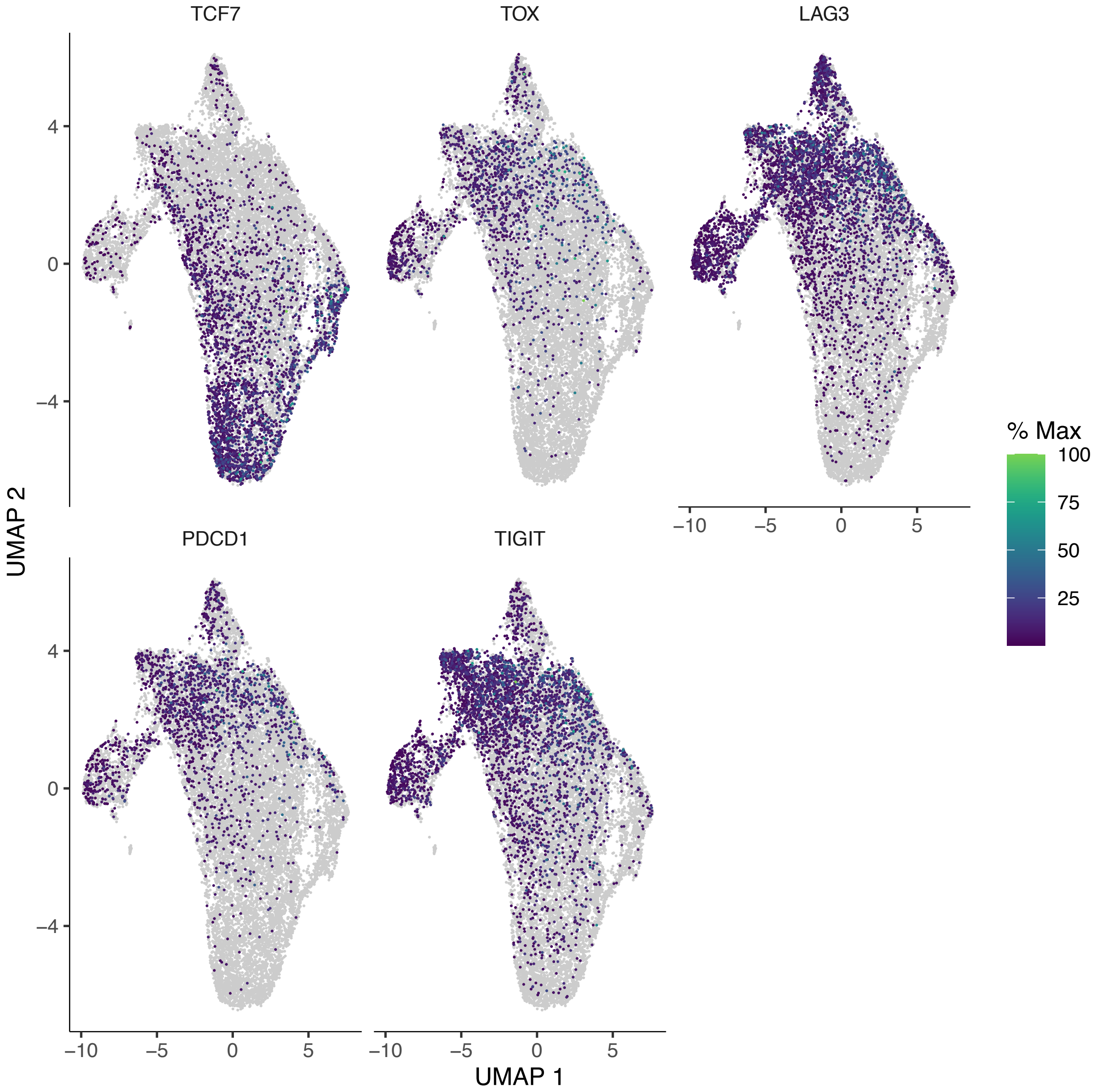


**Figure S3: CD8 T cell UMAP plot – expression of key genes in melanoma dataset**

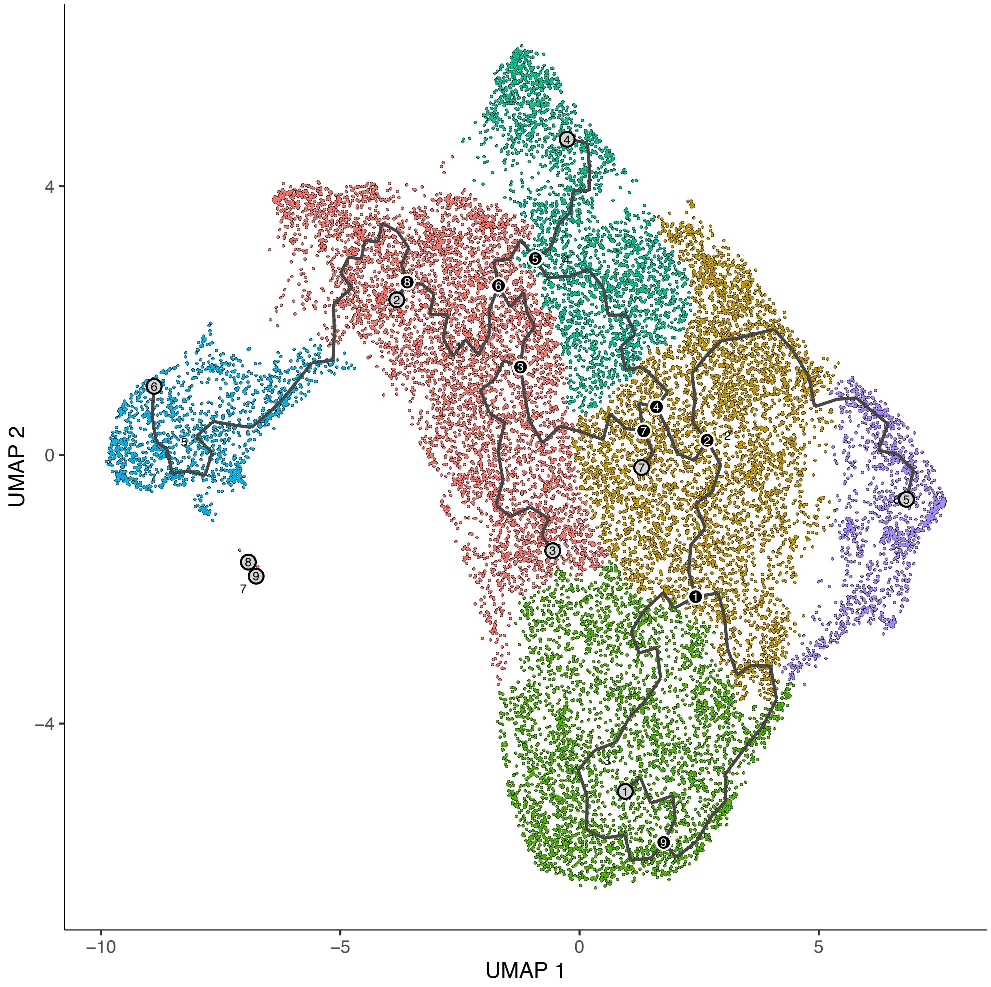


**Figure S4**: part 1 – clusters

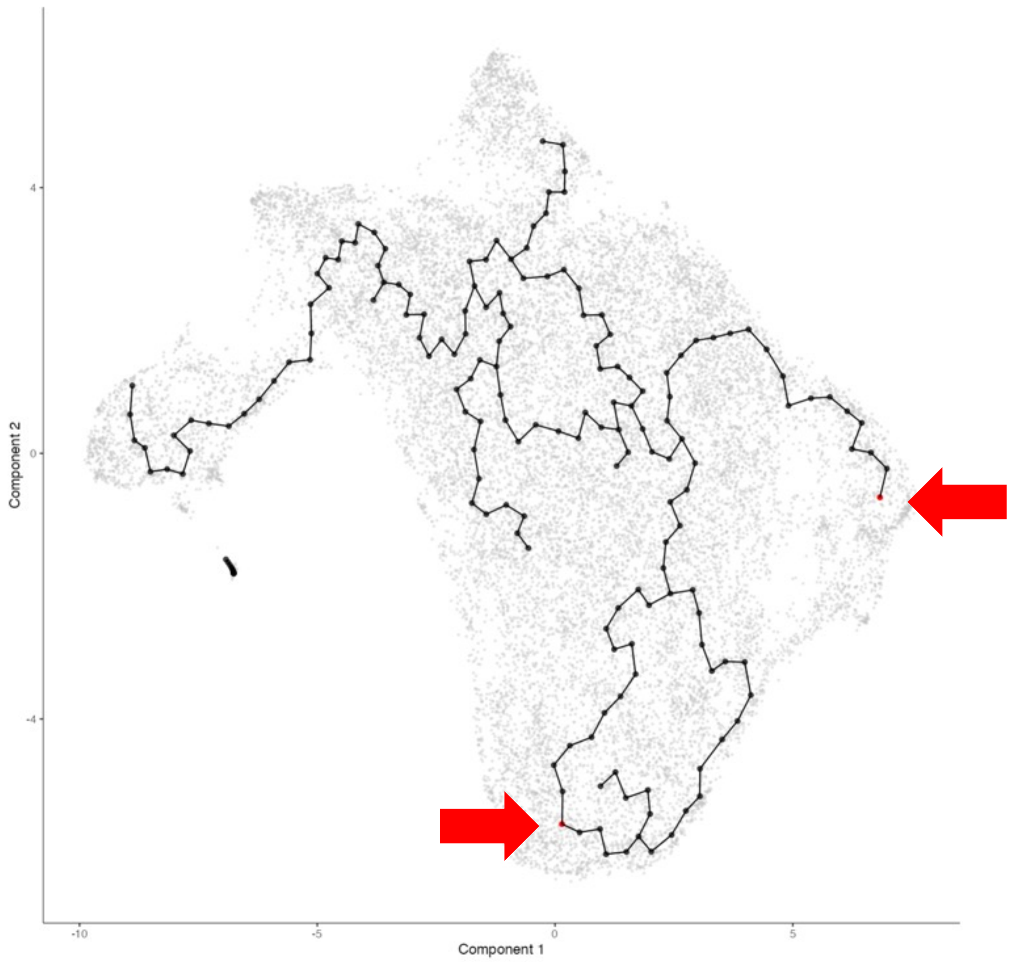


**Figure S4**: **CD8 T cell UMAP plot – Monocle3 principal graph and clusters for melanoma dataset**
top: clusters; bottom: principal graph and root cells
**Parameters used:** Number of principal components: 18; Clustering: k=20, resolution=0.0001

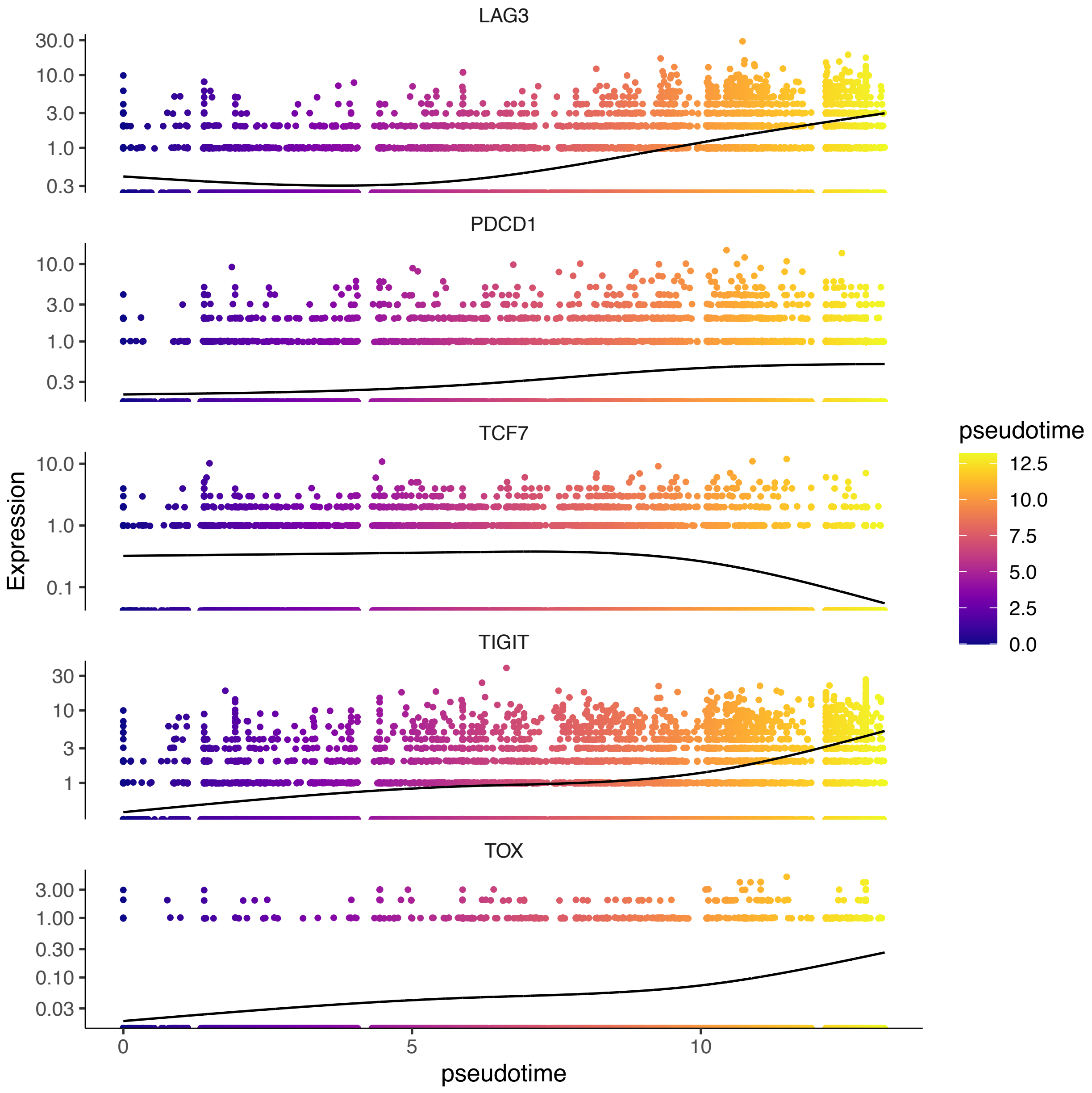


**Figure S5**: **gene expression vs. pseudotime in basal cell carcinoma dataset**
Expression of TCF7, the primary marker of the progenitor exhausted CD8 T cell state, starts high, increases very slightly and then decreases relative to pseudotime. TOX, the primary marker of the terminally exhausted CD8 T cell state, is monotonically increasing across pseudotime. Immune checkpoint genes LAG3, PDCD1, and TIGIT increase across pseudotime (with a slight decrease in LAG3 at the beginning).

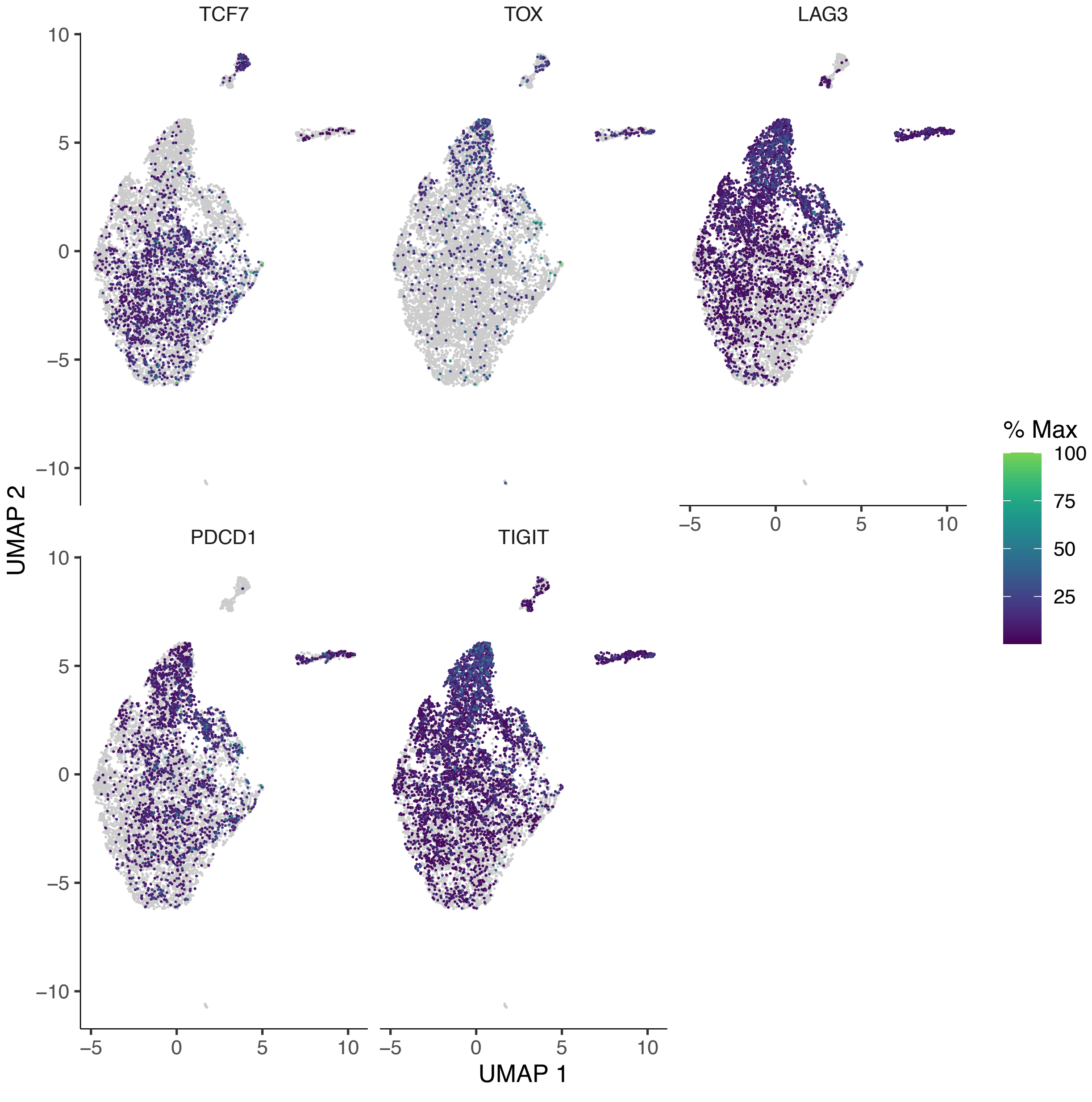


**Figure S6: CD8 T cell UMAP plot – expression of key genes – BCC dataset**

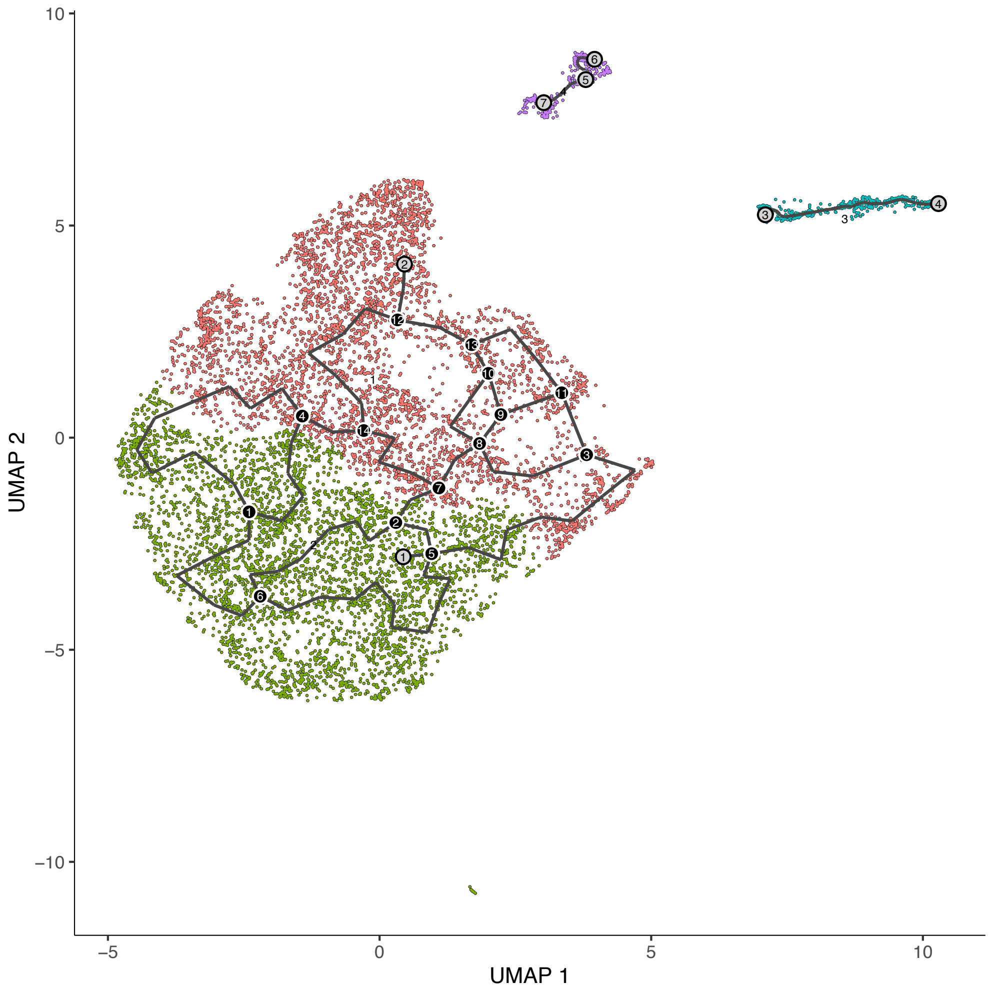


**Figure S7**: part 1 – clusters

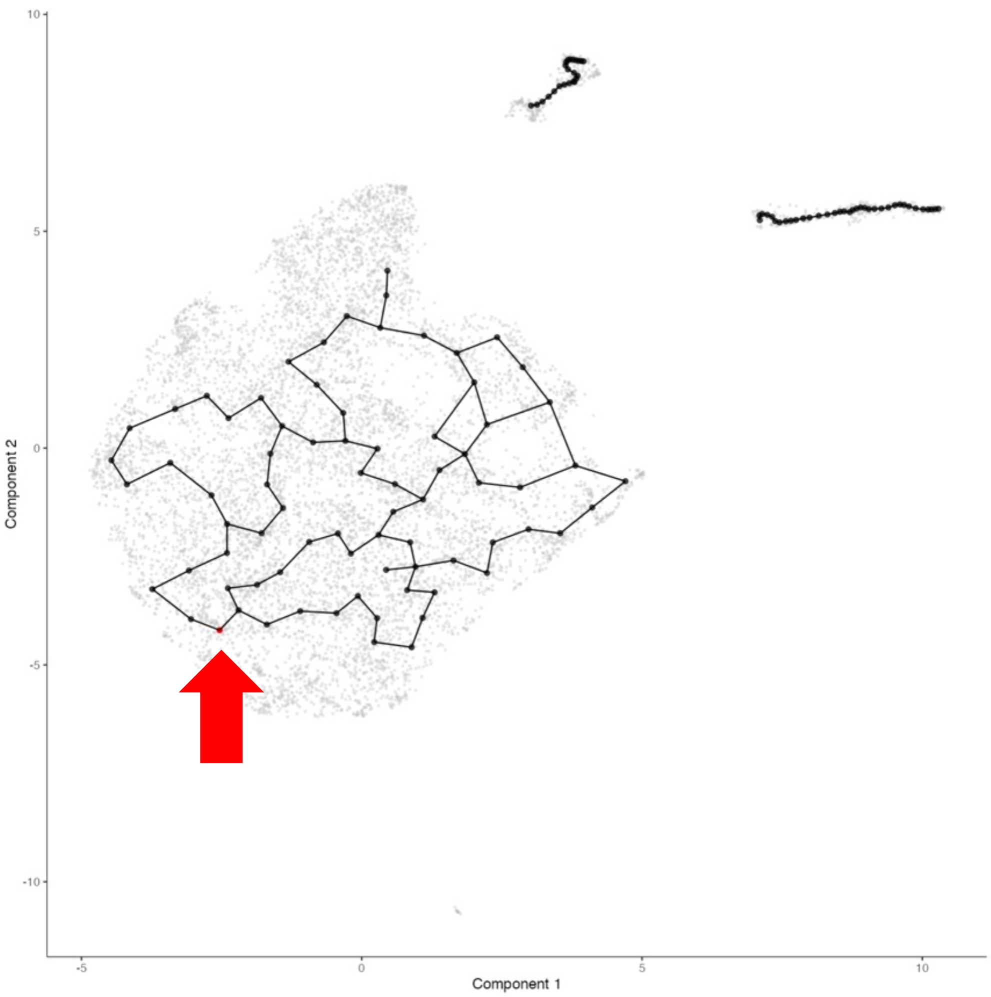


**Figure S7**: **CD8 T cell UMAP plot – Monocle3 principal graph and clusters – BCC dataset**
top: clusters; bottom: principal graph and root cells
**Parameters used:** Number of principal components: 15; Clustering: k=20, resolution=0.0001

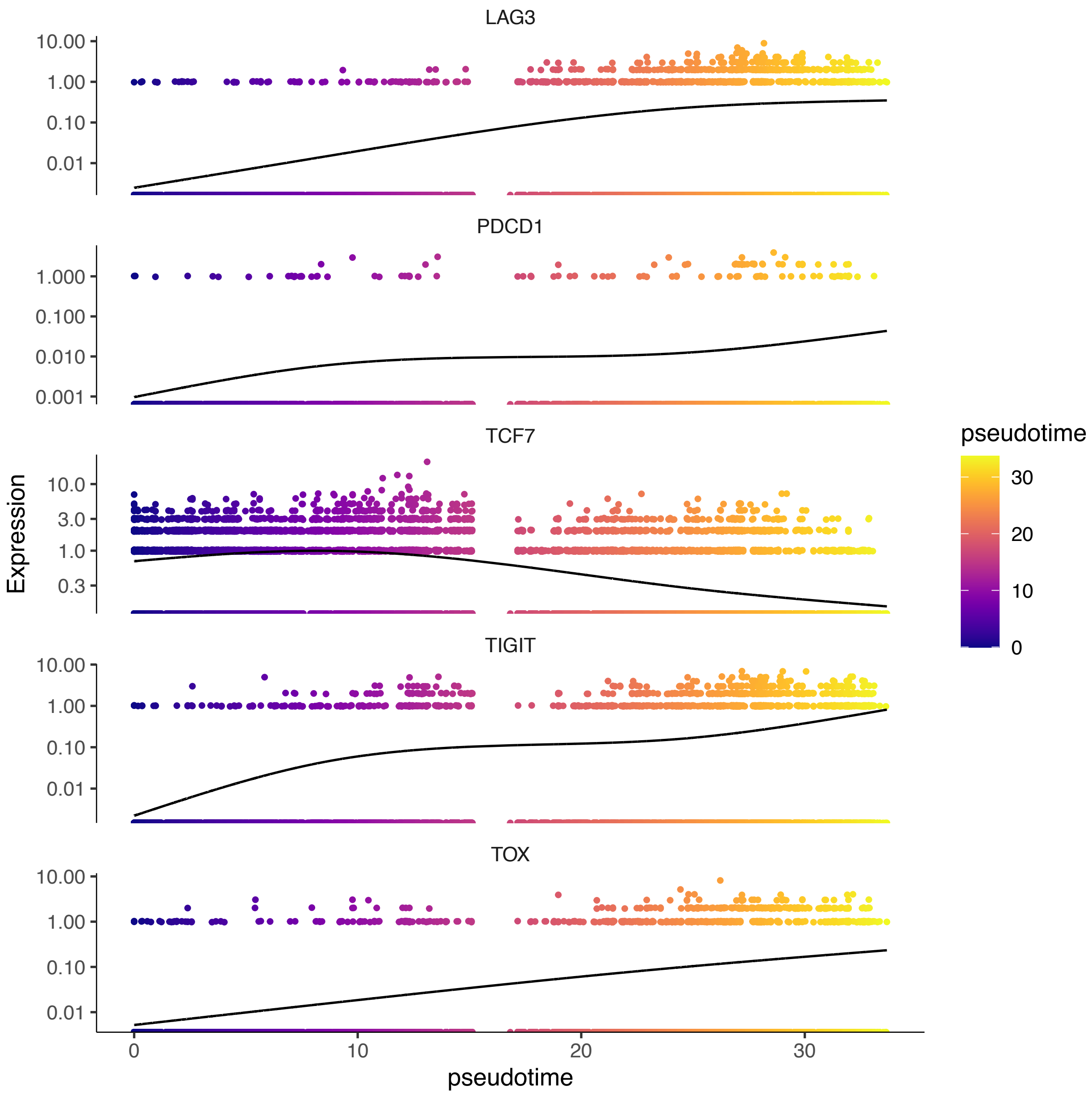


**Figure S8**: **gene expression vs. pseudotime in chronic HIV dataset**
Expression of TCF7, the primary marker of the progenitor exhausted CD8 T cell state, starts high before increasing slightly and then decreasing across pseudotime. TOX, the primary marker of the terminally exhausted CD8 T cell state, is monotonically increasing across pseudotime. Immune checkpoint genes LAG3, PDCD1, and TIGIT are also monotonically increasing across pseudotime.

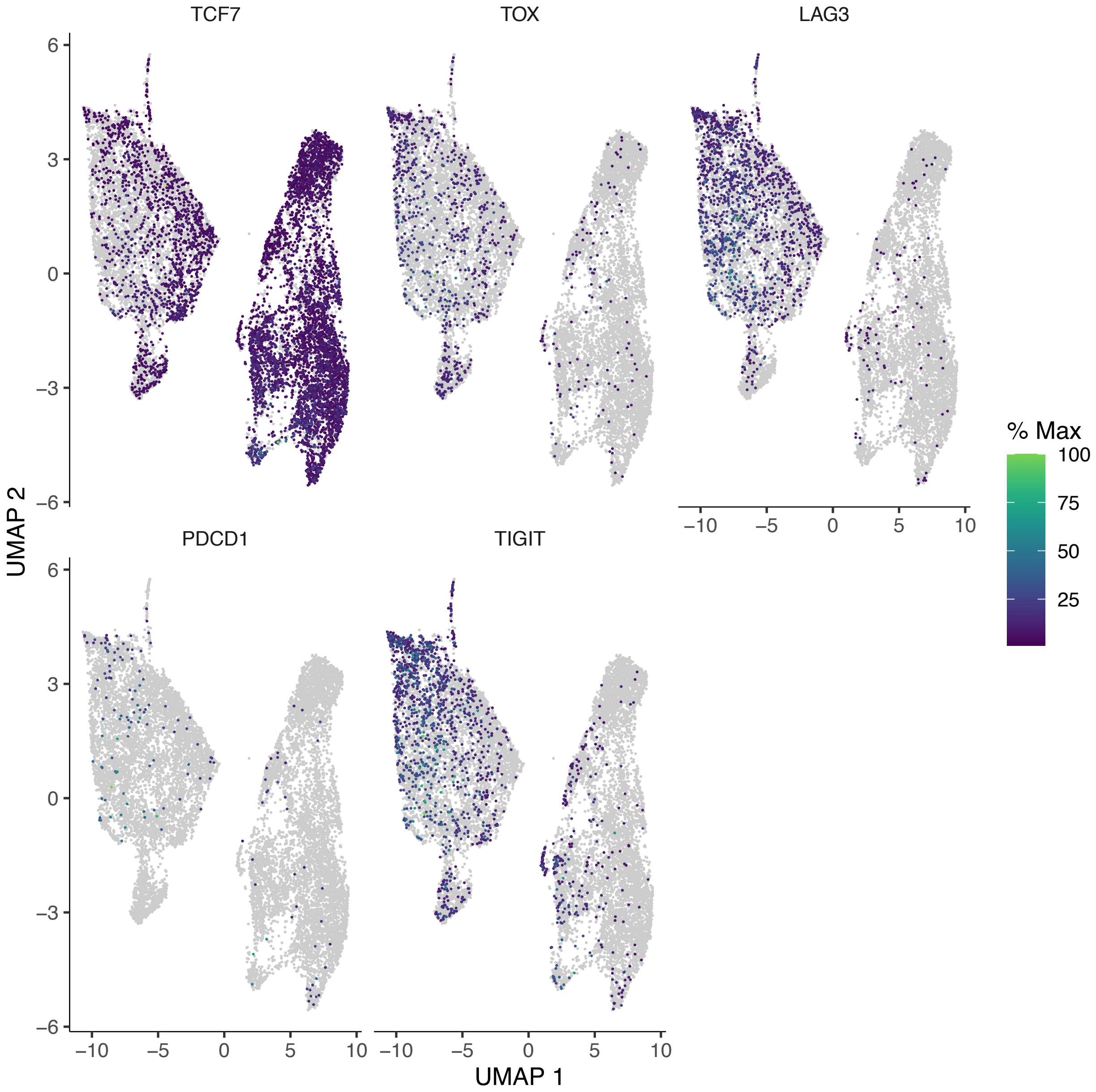


**Figure S9: CD8 T cell UMAP plot – expression of key genes – HIV dataset**

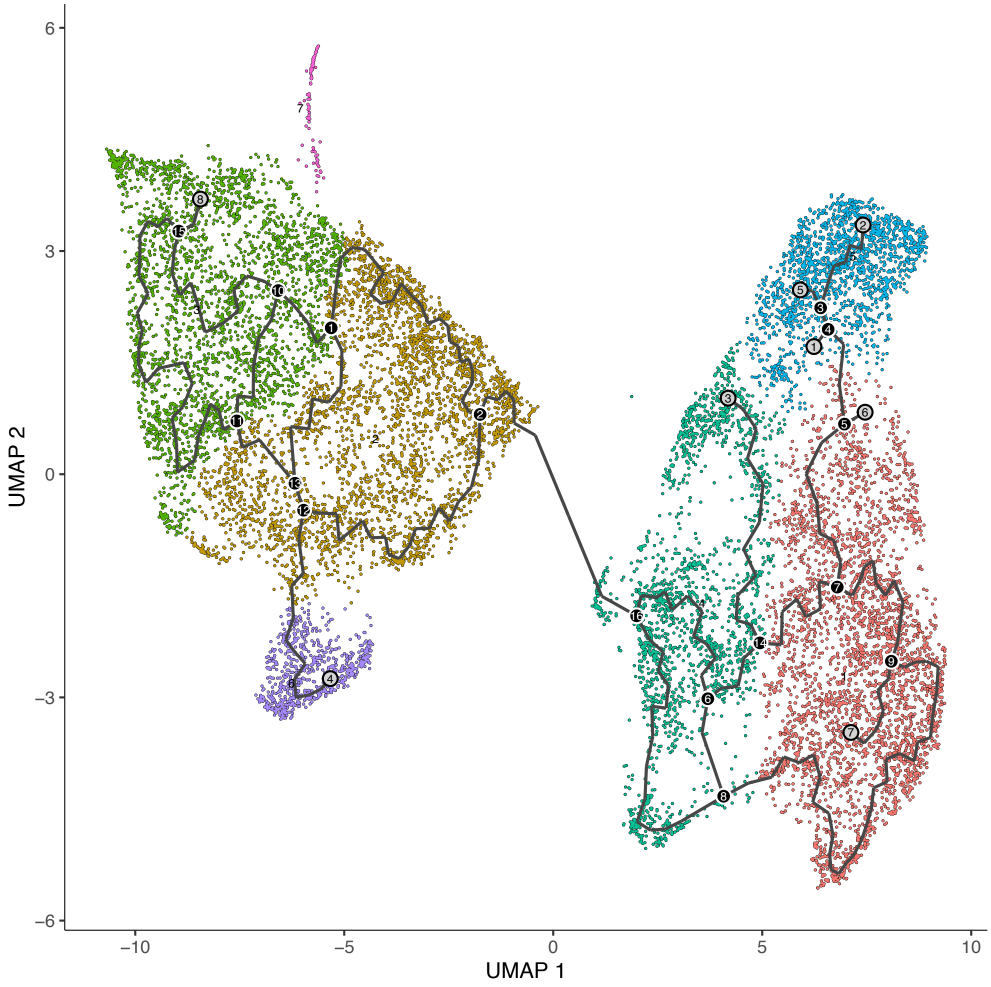


**Figure S10**: part 1 – clusters

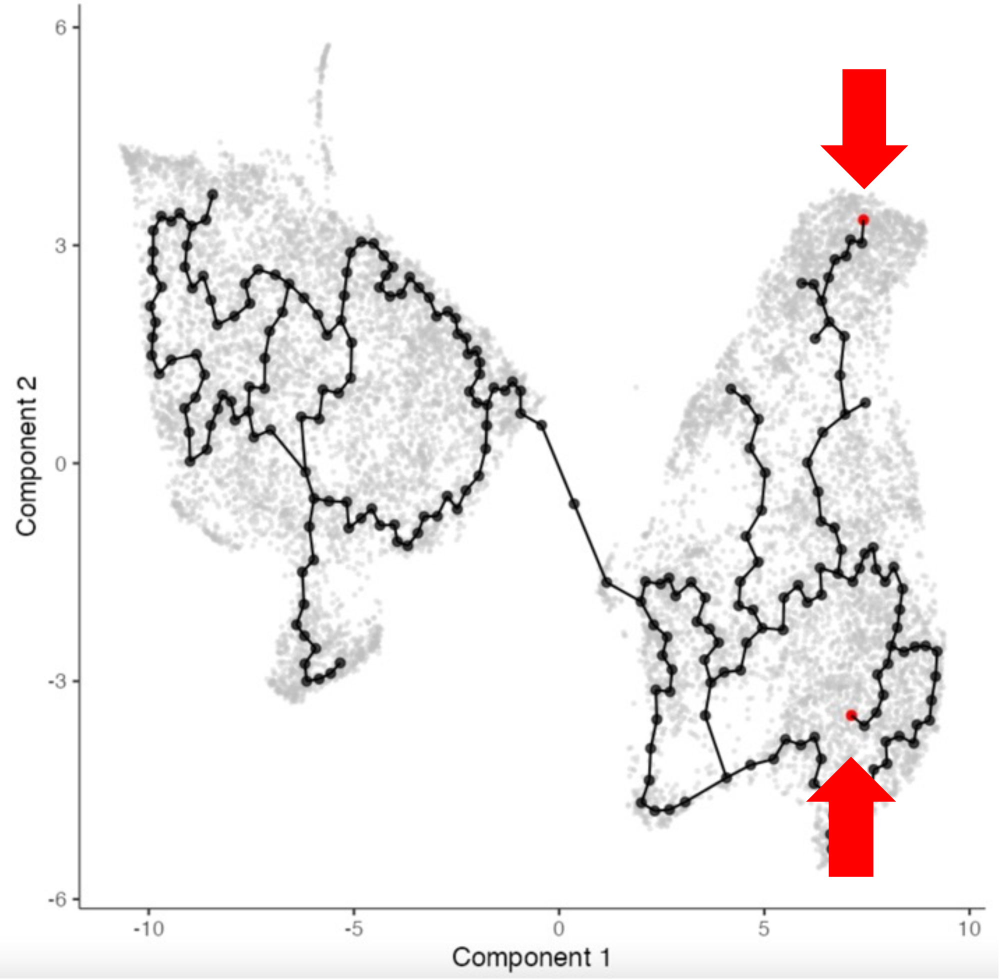


**Figure S10**: **CD8 T cell UMAP plot – Monocle3 graph and clusters – HIV dataset**
top: clusters; bottom: principal graph and root cells
**Parameters used:** Number of principal components: 10; Clustering: k=20, resolution=0.0001; 'learn_graph()': use_partition=FALSE

PATHWAY FIGURES – KEYS:

| | |
|---|---|
| ■ (dark green) | tumor -- melanoma |
| ■ (light green) | tumor -- BCC |
| ■ (salmon) | viral -- HIV |

**Pathway Categories:**

**Immune System**
**Signal Transduction**
**Cell-Cell Communication**
**Responses to Stimuli**
**Other**

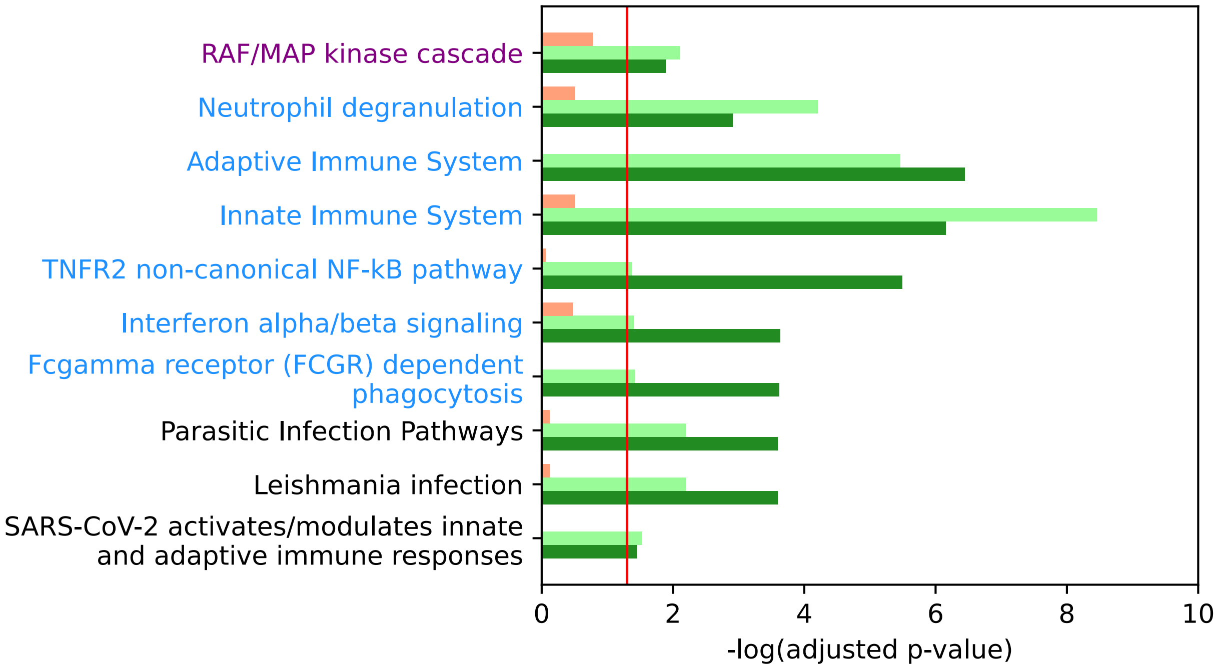


**Figure S11: Pathway enrichment of genes up-regulated in the most exhausted macrophage samples – unabridged, tumor-specific**

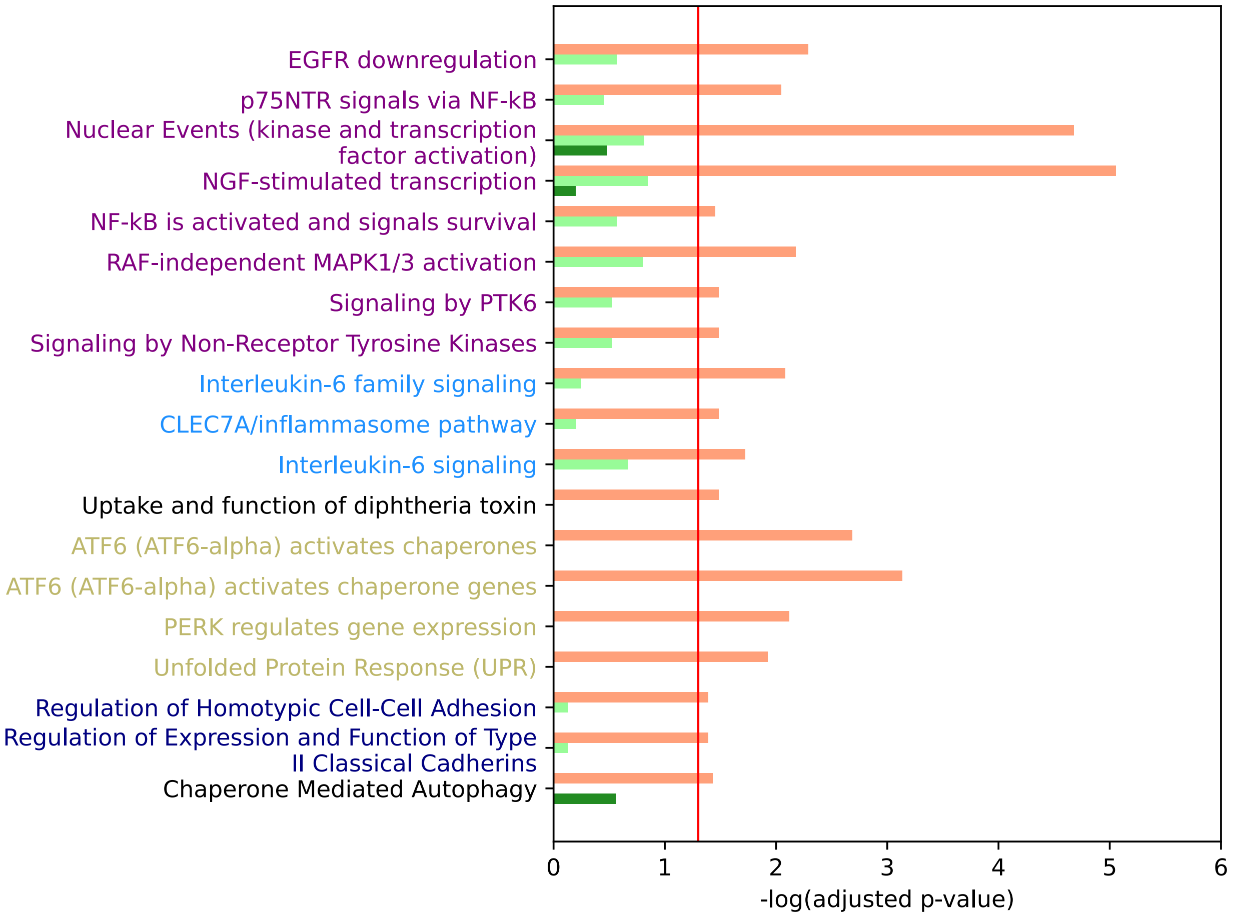


**Figure S12: Pathway enrichment of genes up-regulated in the most exhausted macrophage samples – unabridged, viral-specific**

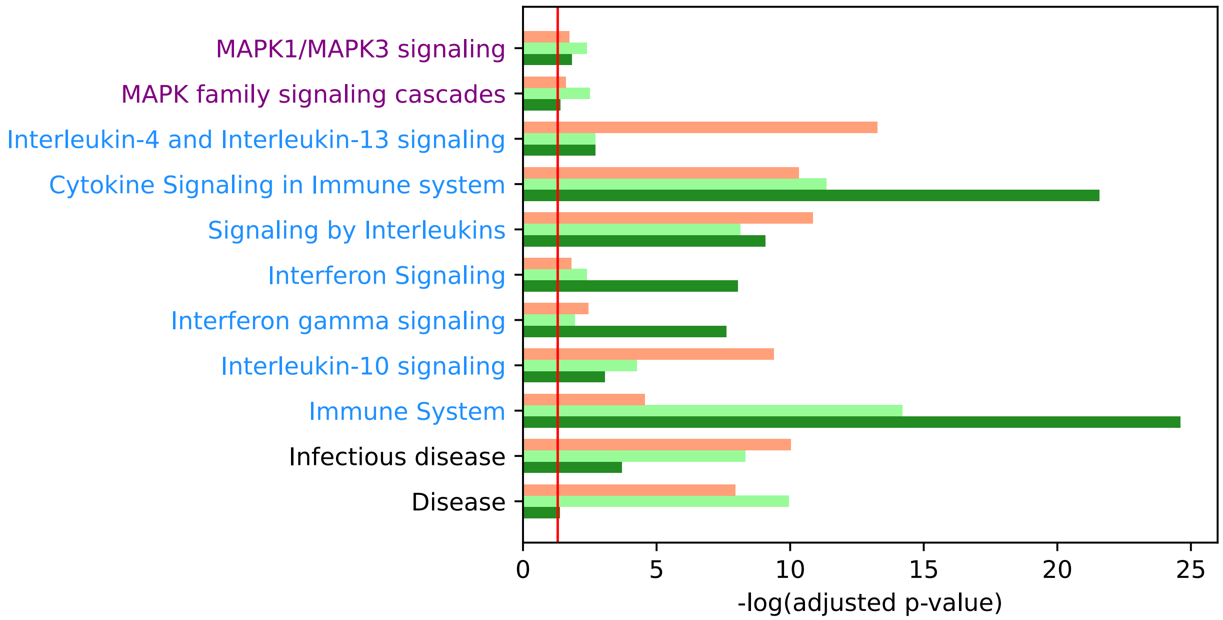


**Figure S13**: **Pathway enrichment of genes up-regulated in the most exhausted macrophage samples – unabridged, shared**

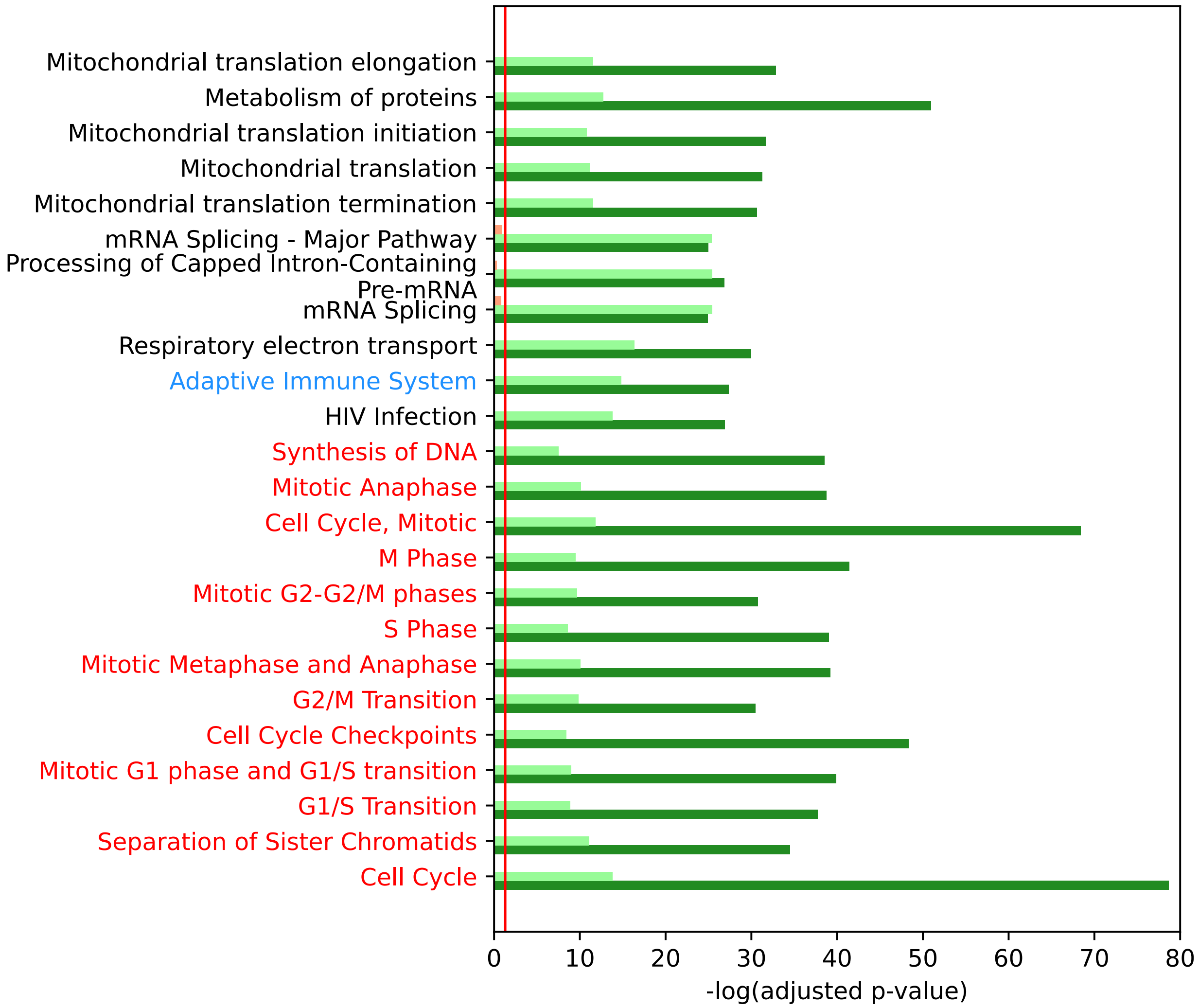


**Figure S14: Pathway enrichment of genes up-regulated in the most exhausted CD8 T cell samples – unabridged, tumor-specific**

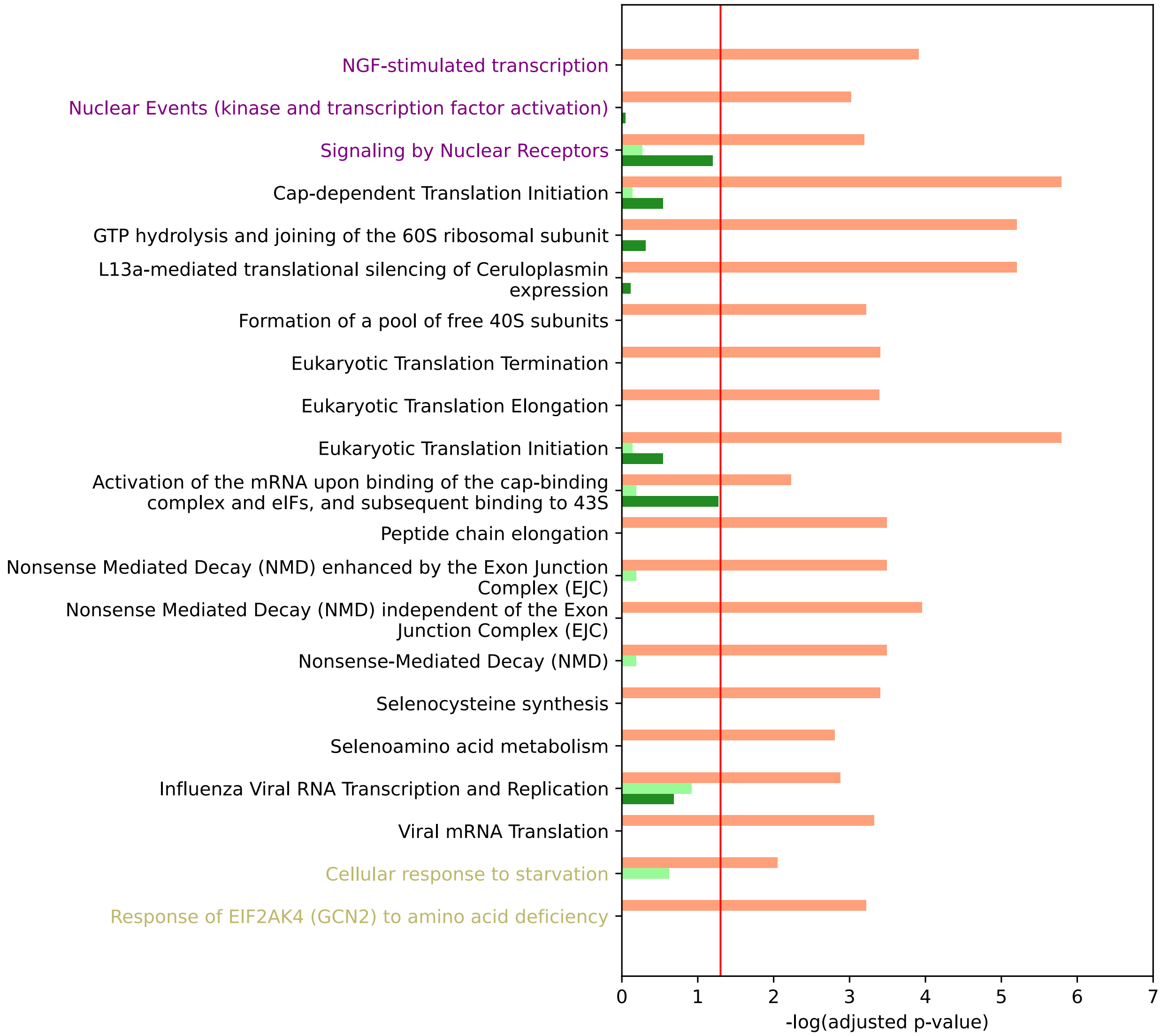


**Figure S15**: **Pathway enrichment of genes up-regulated in the most exhausted CD8 T cell samples – unabridged, viral-specific**

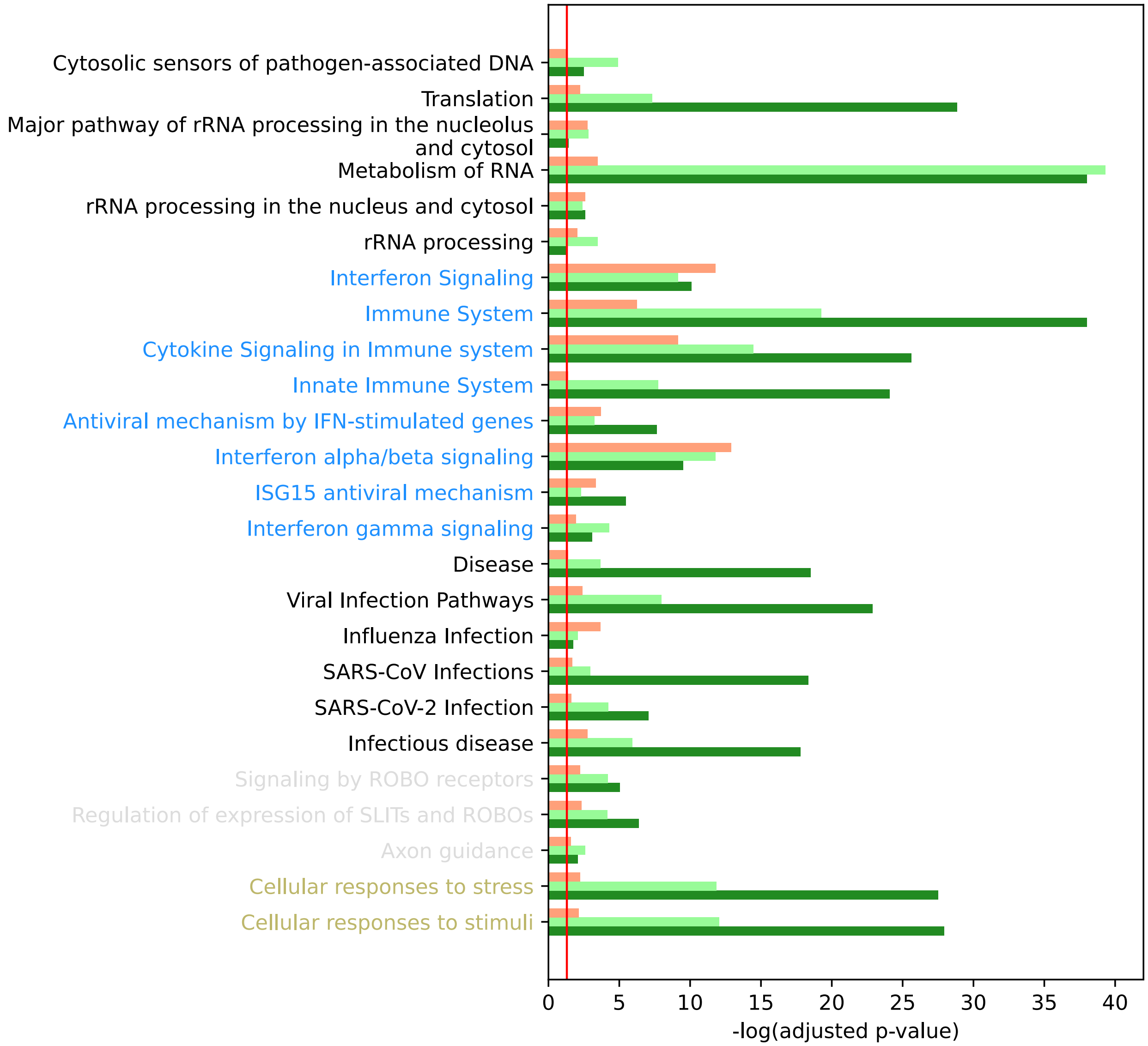


**Figure S16**: **Pathway enrichment of genes up-regulated in the most exhausted CD8 T cell samples – unabridged, shared**

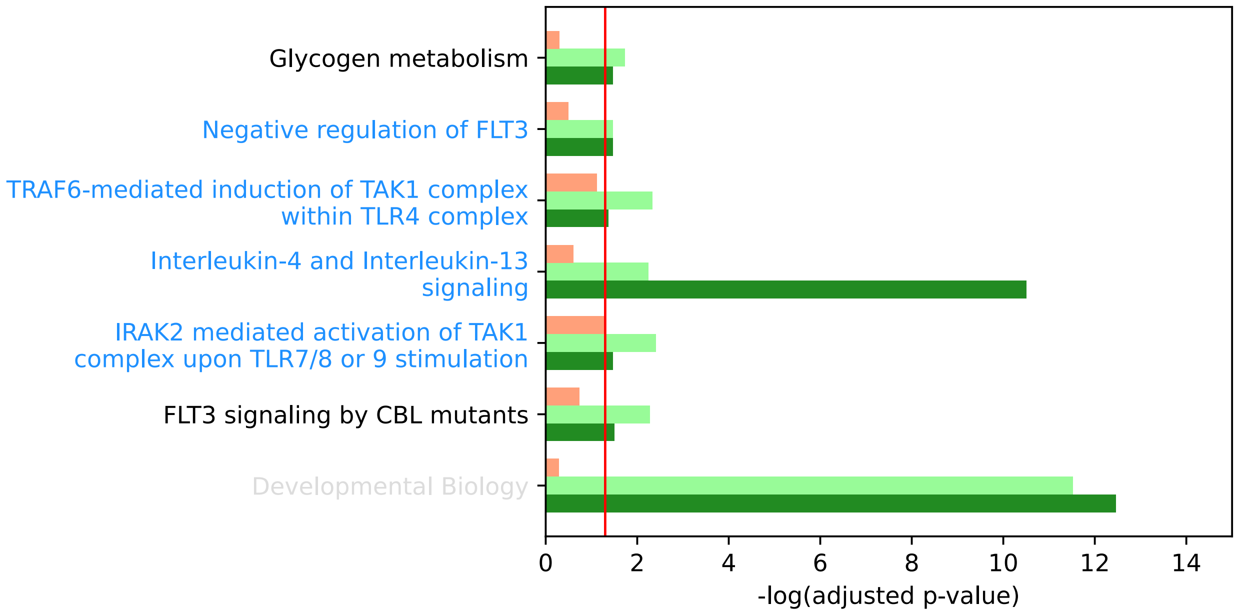


**Figure S17**: **Pathway enrichment of genes down-regulated in the most exhausted CD8 T cell samples – unabridged, tumor-specific**

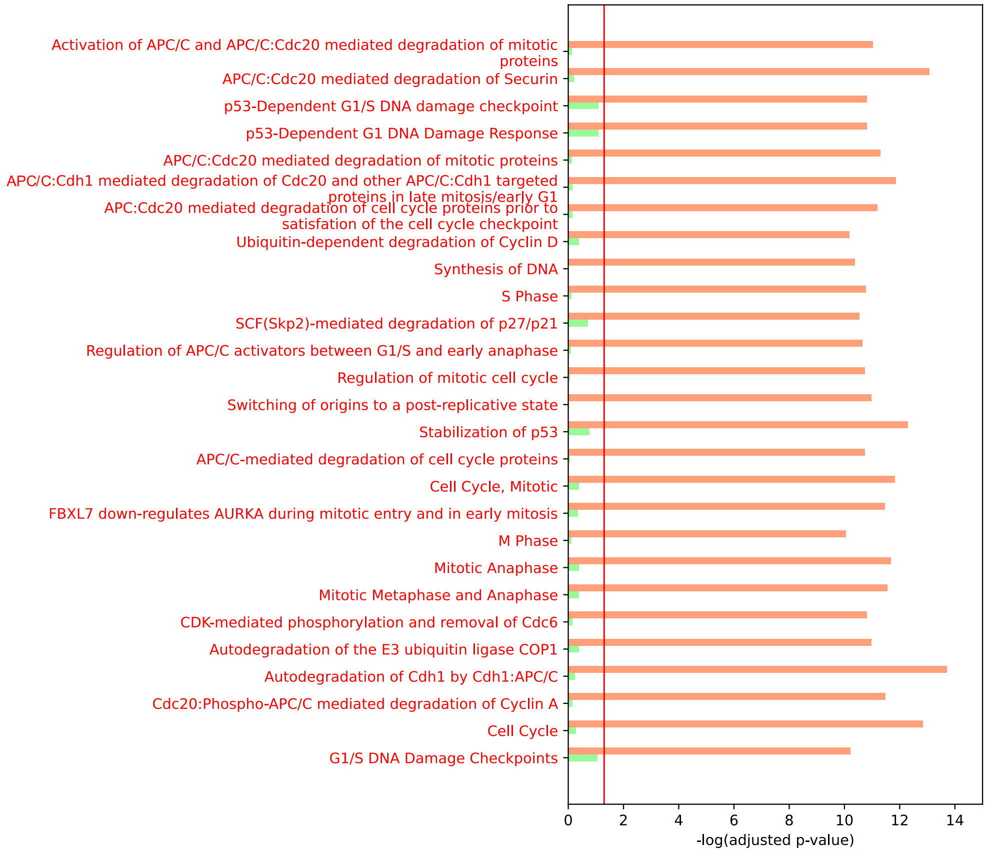


**(part 1 of 3)**

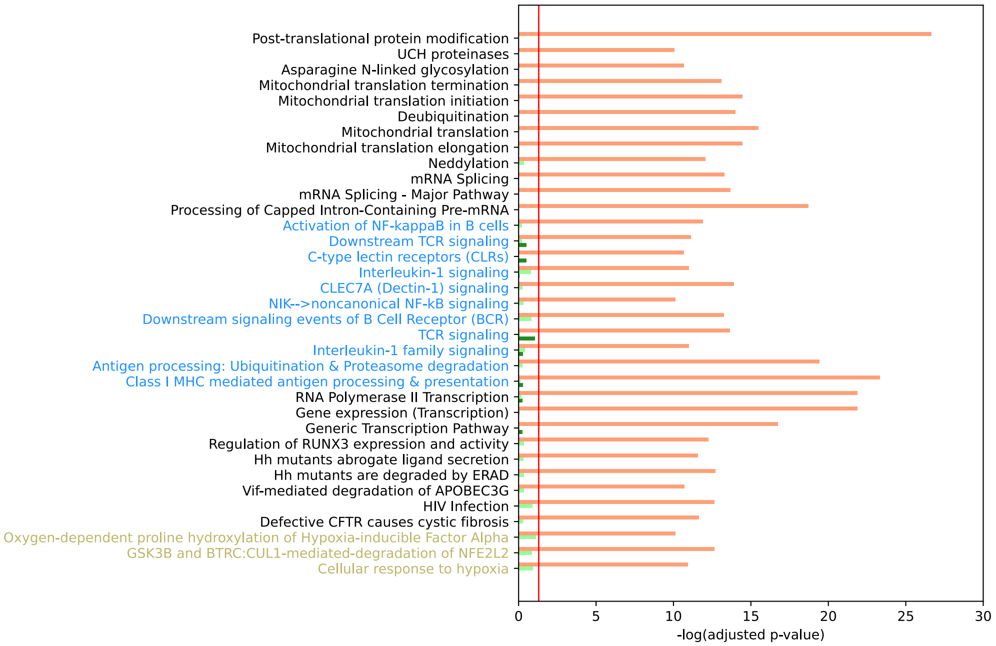


**(part 2 of 3)**

**Figure S18: Pathway enrichment of genes down-regulated in the most exhausted CD8 T cell samples – unabridged, viral-specific**

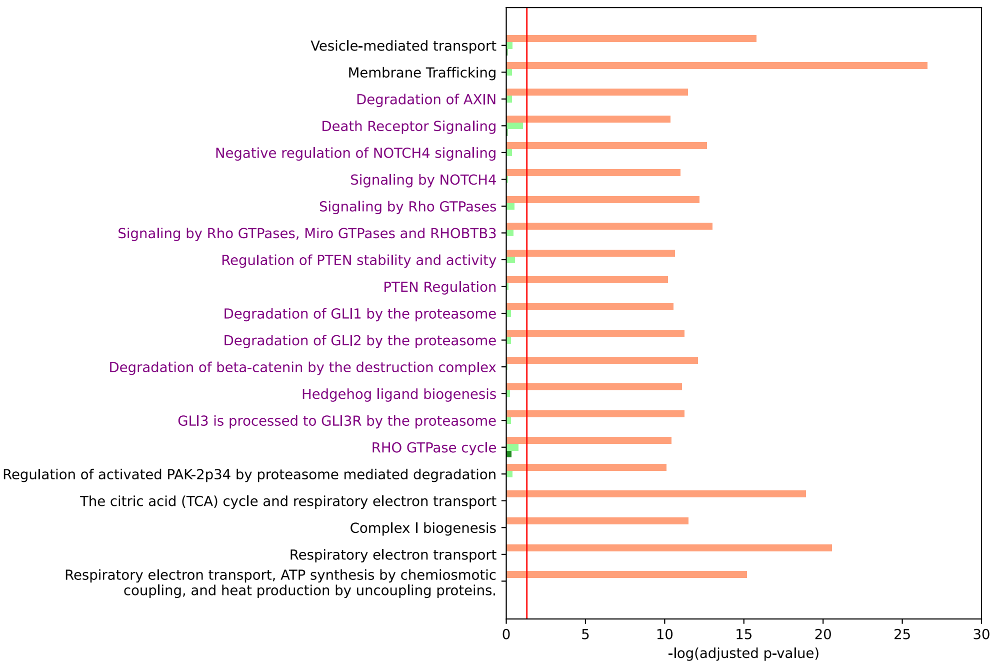


**(part 3 of 3)**

**Figure S18: Pathway enrichment of genes down-regulated in the most exhausted CD8 T cell samples – unabridged, viral-specific**

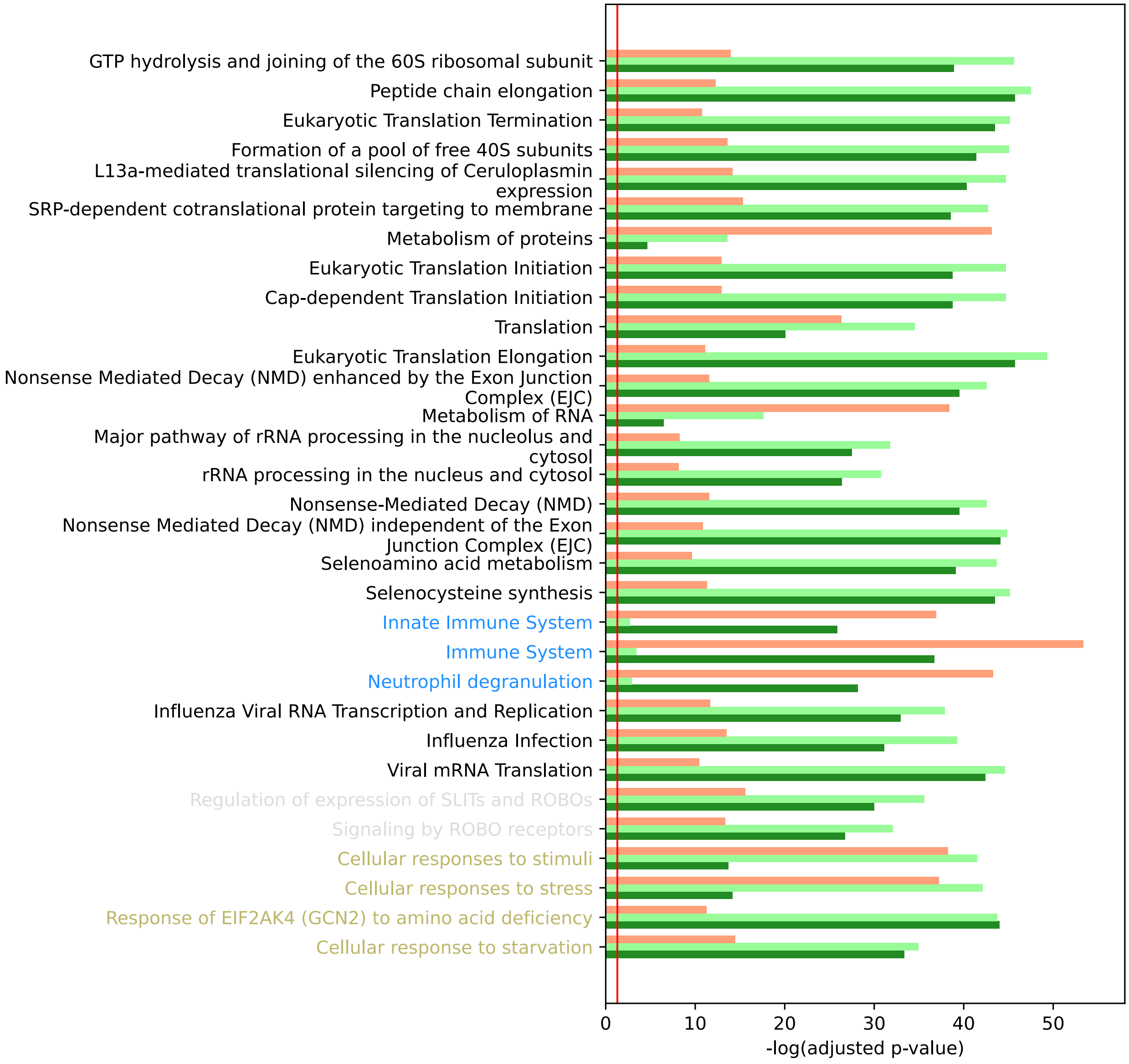


**Figure S19: Pathway enrichment of genes down-regulated in the most exhausted CD8 T cell samples – unabridged, shared**

| | Dataset | | |
|---|---|---|---|
| **cell type** | Li et al., 2019 | Yost et al., 2019 | Wang et al., 2020 |
| CD8 T cells | 19741 | 8136 | 15202 |
| macrophages | 5298 | 1144 | 1895 |
| natural killer cells | 5523 | 196 | - |
| B cells | 3999 | 111 | 4440 |
| plasma cells | 1283 | 1710 | 404 |
| CD4 T cells | - | 3619 | - |
| T regulatory cells | - | 2039 | - |
| tumor cells | - | 2492 | - |
| cancer-associated fibroblasts | - | 626 | - |
| dendritic cells | - | 396 | - |
| endothelial cells | - | 235 | - |
| melanocytes | - | 88 | - |
| myofibroblasts | - | 162 | - |
| unknown | 1717 | - | - |
| total | 37561 | 20954 | 21941 |

**Table S1: Cell counts by cell type for each of three datasets**
